# A genome-wide genetic interaction platform for MRSA reveals connections between cell division and the cell envelope

**DOI:** 10.64898/2026.08.26.747279

**Authors:** Sumana Bhowmick, Félix Ramos-León, Kumaran S. Ramamurthi, Seth W. Dickey

## Abstract

The bacterial cell cycle is an ensemble of integrated pathways that coordinates cell growth and division. Although these pathways contain a suite of promising antimicrobial targets, their functional organization and interrelationships have only been sparsely mapped. Here, we established a dual-CRISPRi platform for systematic genetic interaction profiling in methicillin-resistant *Staphylococcus aureus* (MRSA), a major drug-resistant pathogen. Our platform achieved genome-wide coverage across a selected set of 51 cell cycle genes, surveying 115,497 gene-pairs and uncovering hundreds of genetic interactions, both positive (suppressors) and negative (synthetic sick/lethal). These interactions identified two major connections between cell division and the cell envelope. First, we identified a suppressive relationship between a defined set of cell division genes, including components of the major septal peptidoglycan biosynthesis complex PBP1-FtsWL-DivIBC, and fatty acid or phospholipid biosynthesis. Simultaneous inhibition of these lipid biosynthesis pathways suppressed the fitness and morphological defects caused by knocking down expression of cell division genes, revealing that toxic membrane accumulation contributes to the lethality of disrupting cell division. Second, we uncovered a synthetic lethal relationship between a distinct set of cell division genes and the conserved *dltXABCD* operon, linking cell division to the modification of teichoic acid surface polymers. Our results establish a versatile platform for interrogating selected gene sets in MRSA, providing a map of cell cycle interactions, and uncovering links between cell division and cell envelope pathways.

## Introduction

Bacterial physiology is organized through an interconnected network of cellular pathways. For the cell cycle, these include DNA replication, chromosome segregation, macromolecular biosynthesis, septum formation, cell division, and daughter cell splitting^1–3^. These events are tightly coupled, and functional redundancy is common, helping to ensure faithful execution under myriad environments and stresses^2–4^. Moreover, because many of the factors acting within these pathways are essential, the cell cycle contains a suite of promising targets for the development of new antibiotics and antimicrobials.

Despite a longstanding interest in understanding the cell cycle, however, the functional organization among its pathways remains incompletely mapped. In addition, bacteria exhibit substantial variation in the mechanisms and proteins that execute the cell cycle, reflecting ancient evolutionary divergence as well as underlying differences in physiology and cell morphology^5,6^. This limits the generalizability from traditional model systems and motivates direct investigation in drug-resistant pathogens^7–9^. Methicillin-resistant *Staphylococcus aureus* (MRSA) is a prevalent and leading cause of antibiotic-resistant infections globally and, ranks among the deadliest pathogens^10,11^. With no vaccine available, maintaining and expanding antimicrobial options remain key to reducing the burden of disease caused by MRSA.

Surveying for genetic interactions is a powerful tool that has revealed underlying pathway architecture^12,13^. These interactions arise from gene products that function within essential, parallel, or regulatory pathways and manifest experimentally when the disruption of two genes yields a fitness that deviates from the expected outcome based individual gene perturbations. Such interactions can be negative when the gene pair is synthetic sick or lethal, and positive when one perturbation compensates for the fitness defect of another. Existing large scale genetic interaction profiling studies in bacteria have leveraged model and genetically tractable bacterial species^14–17^, revealing how cell envelope synthesis and stress response systems interface with cell growth and division. While case-by-case studies including genetic interactions have revealed insight into MRSA physiology and virulence^18–22^, we lack a systematic and scalable approach for *S. aureus* in part due to the low efficiency of natural competence and robust restriction-modification barriers^23–25^.

Here, we report the development, validation, and application of a dual-CRISPRi platform for genome-scale genetic interaction mapping in *S. aureus*, overcoming technical hurdles to build large heterogenous libraries capable of simultaneously silencing two genes within a cell. We applied our platform to cell cycle pathways and, after surveying 115,497 possible gene–gene combinations, we identified hundreds of genetic interactions and validated two major clusters that uncover connections between cell division proteins and the cell envelope.

## Results

### Dual-CRISPRi platform for systematic genetic interaction mapping in S. aureus

To implement genome-scale genetic interaction screening in *S. aureus*, we adopted the well-established CRISPR-interference (CRISPRi) gene-silencing approach^26–28^, which is amenable to creating high-diversity single-guide (sg)RNA libraries of designed spacer sequences and evaluating changes in spacer frequency within a heterogenous cell population to measure abundance-based relative fitness^15,16,29^. We engineered the MRSA strain USA300 JE2^30^ for inducible expression of a chromosomally integrated *Streptococcus pyogenes* catalytically-dead *cas9* (*dcas9)* under control of a P_tet_ promoter variant optimized to minimize leaky expression^31,32^.

We integrated this cassette into the *hsdR* locus that encodes the type I restriction nuclease found widely across *S. aureus* strains, thereby disrupting the gene and increasing transformation efficiency by more than 100-fold^33^ (**Fig. 1a, b**). We then developed a dual-sgRNA delivery shuttle plasmid, pSD14, carrying two orthogonal Golden Gate assembly sites (BsaI and BbsI) for generation of two sgRNAs under control of constitutive promoters^34,35^. We positioned the antibiotic resistance marker (*cat*) between the sgRNA scaffolds to prevent recombinational loss of the sgRNA at the BbsI assembly site (**Fig. 1c**). Finally, we inserted a BsmBI Golden Gate assembly site to accommodate DNA barcodes for marking individual sgRNAs at the BbsI site. This enabled identification and quantification of sgRNA pairs via paired-end amplicon sequencing of the BsaI sgRNA and BsmBI barcode. Together, JE2 *hsdR*::*dcas9-tetR* and pSD14 constitute a platform for quantifying fitness consequences of pairwise gene knockdowns across the *S. aureus* genome. *Validation of the Dual-CRISPRi in S. aureus platform*

**Fig. 1.**
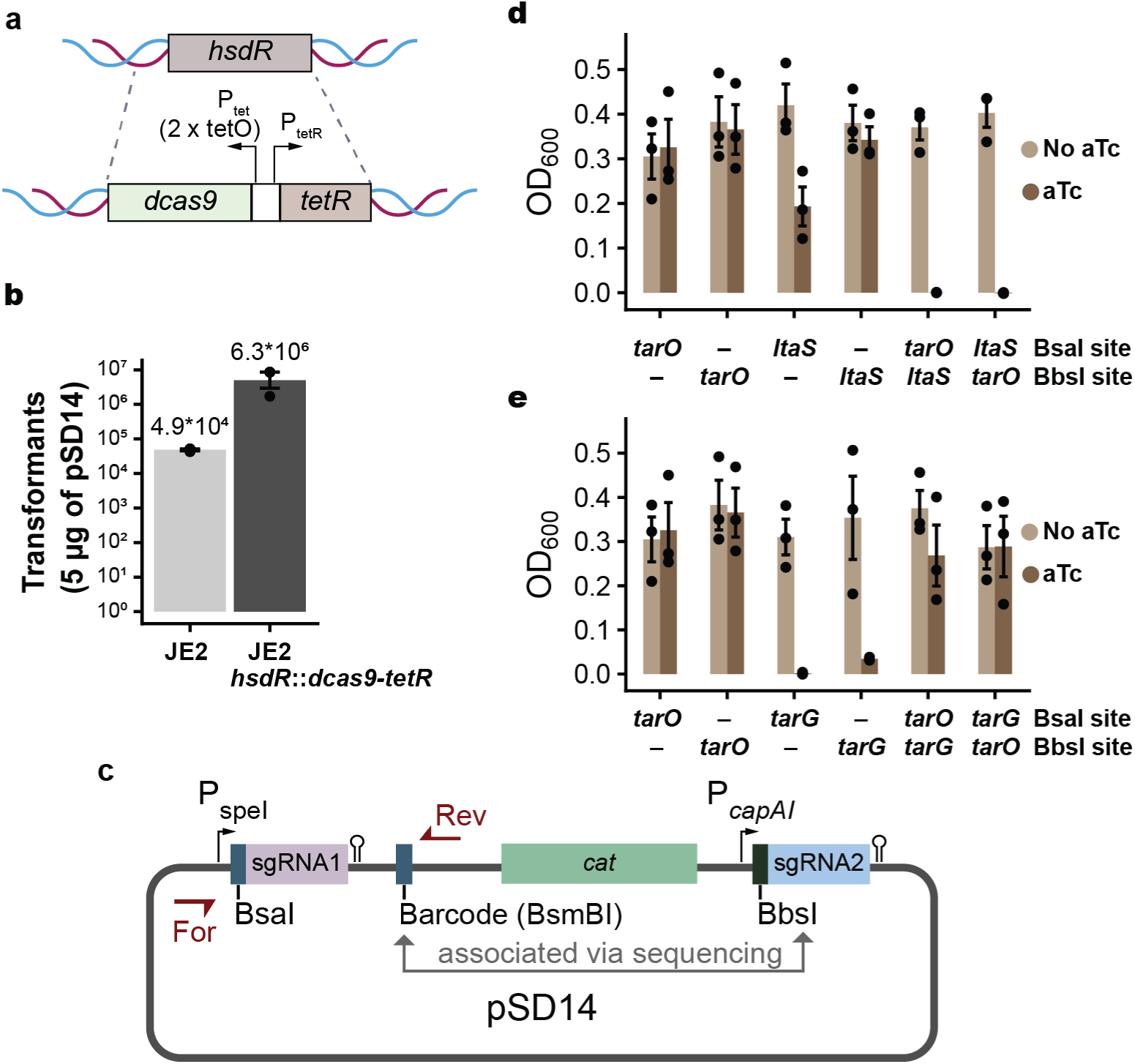
A high-throughput dual-CRISPRi platform in *S. aureus* for uncovering genetic interactions. **a.** Inducible *dcas9* expression cassette chromosomally integrated into *S. aureus* JE2 replacing *hsdR,* which encodes a restriction enzyme. **b.** Comparison of transformation efficiency of pSD14 Dual-sgRNA delivery vector between the engineered JE2 *hsdR*::*dcas9-tetR* strain and the JE2 strain competent cells; (n=3 independent replicates, p < 0.01). **c.** Schematic of the pSD14 dual sgRNA delivery vector. Spacers are assembled at the BsaI or BbsI sites. Barcodes are assembled at the BsmBI site. Amplicon priming sites are denoted by R1 and R2. **d, e.** Growth of JE2 *hsdR*::*dcas9-tetR* in presence and absence of the inducer expressing paired sgRNAs targeting *tarO* and *ltaS* (**d**) or *tarO* and *tarG* (**e**). n = 3 biological replicates; bars show mean ± standard error; for samples with aTc, p < 0.001 using ANOVA followed by Tukey’s HSD test comparing (**d**) dual targeting of *tarO* and *ltaS* with both single-targeting groups and (**e**) single targeting of *tarG* with single targeting of *tarO* or dual targeting of *tarO* and *tarG*.

We validated the platform by first targeting known genetic interactions in *S. aureus*. Combining *tarO* and *ltaS* loss-of-function mutations results in a negative (synthetic lethal) interaction due to the simultaneous disruption of wall teichoic acid (WTA) and lipoteichoic acid (LTA) biosynthetic pathways^18,36^ In contrast, disrupting *tarO* function exhibits a positive (suppressive) interaction with the loss of *tarG* function^19,20^. Our dual-CRISPRi platform recapitulated both interactions in a *dcas9*-dependent manner (**Fig. 1d, e**), thereby demonstrating the ability to detect negative and positive genetic interactions.

We next constructed a pilot-scale dual-sgRNA library that included the known genetic interactions with *tarO* (**Fig. 2a**). In total, we paired 6 sgRNAs at the BbsI site (4 targets plus 2 non-targeting controls [NTC]) with 50 sgRNAs at the BsaI sites (2 independent spacers for each of 24 targets plus 2 NTCs). We recovered ∼7×10^8^ transformants and ∼2×10^5^ transformants, respectively (>600 × coverage) after transforming the library serially to *E. coli* DC10B and *S. aureus* JE2 *hsdR*::*dcas9-tetR*. After growth in parallel cultures with or without supplementation of the inducer aTc followed by amplicon sequencing (**Fig. 2b**), we calculated the relative fitness (adjusted log_2_(fold-change), see Methods) for single-gene silencing (W_i_ and W_j_ at the BsaI and BbsI positions, respectively) and dual-gene silencing (W_ij_). We then used the Product model^13,29^ to derive genetic interaction scores (*ε_ixj_*). All metrics were well distributed and highly reproducible across replicates (**Fig. 2c and Fig. S1**). Importantly, our platform reproduced interactions with *tarO,* exhibiting a negative interaction with *ltaS* and a positive interaction with *tarG* (**Fig. 2d,e**). Unwittingly, we also reproduced a recently reported interaction between *ltaS* and *cozEb*^37^ (**Fig. 2d**). These data therefore validated end-to-end performance and established that our dual-CRISPRi platform can resolve biologically meaningful genetic interactions in MRSA.

**Figure 2.**
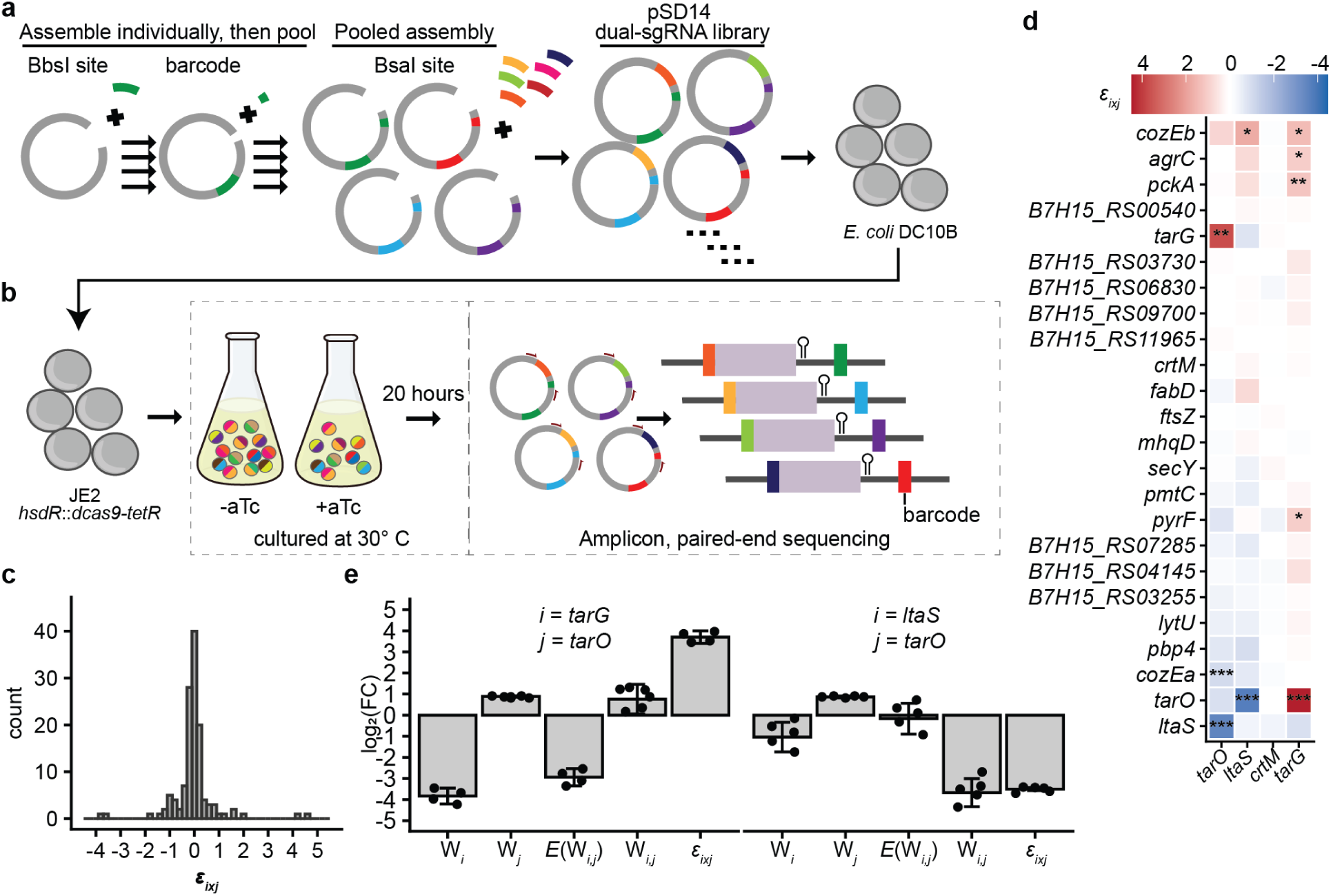
Pilot-scale validation detecting genetic interactions from pooled libraries. **a.** Schematic of the dual-sgRNA library preparation into pSD14. Individual spacer and barcode pairs were assembled at the BbsI and BsmBI sites, respectively. A pool of spacers were assembled at the BsaI site. **b.** The dual-sgRNA plasmid library was transferred to JE2 *hsdR*::*dcas9-tetR* and cells were grown in the presence or abscence of aTc. Plasmid was recovered and amplicons encompassing the BsaI spacer and the barcode were quantified by pair-end sequencing. **c.** Histogram of gene-level (mean, n = 3 biological replicates) genetic interaction (*ε_ixj_*) values. **d.** Heat map of genetic interactions (*ε_ixj_*) (*, fdr < 0.05; **, fdr < 0.01; ***, fdr < 0.001). **e.** Gene-level fitness (W_i_, W_j_, and W_ij_), expected fitness (*E*(W_ij_)) and interaction (*ε_ixj_*) scores recapitulate the known positive (left, i = *tarG*, j = *tarO*) and negative (right, i = *ltaS*, j = *tarO*) interactions.

### Genome-wide survey of genetic interactions with S. aureus cell cycle genes

Building on the success of the pilot library, we focused on elucidating genome-wide interactions of cell-cycle pathways. In the pSD14 BbsI site, we individually assembled and pooled 101 barcoded sgRNAs targeting 51 distinct genes with established roles in DNA replication, chromosome segregation, septation, and cell wall biogenesis. We paired this with a custom designed genome-wide spacer library of 5,674 spacers assembled at the BsaI site. Notably, our genome-wide spacer library was optimized to avoid spacers with predicted off-target binding and it contained two independent spacers for 96% of protein coding sequences (CDSs), with at least one spacer targeting 99% of CDSs. We chose to target all CDSs instead of pursuing an operon approach because 1) promoters may be present within operons^38^ and transcription start sites can be distant from annotated CDSs^39^, 2) findings consistent across CDSs within an operon provide confidence against off-target effects, and 3) the pattern of interactions across CDSs within an operon can be leveraged to generate hypotheses for testing and deconvoluting interacting partners.

We also included 9 and 409 NTCs within the cell-cycle (BbsI, 8.2%) and the genome-wide (BsaI, 7.2%) libraries, respectively, to enable simultaneous parsing of fitness effects for single-gene and dual-gene silencing. After conducting two independent replicates, maintaining >100-fold library coverage in cell numbers and sequencing reads (**Tables S1, S2**), and applying quality and minimum read thresholds to paired-end amplicon sequencing, we observed a distribution of single-gene adjusted fitness values ranging from -6.2 to 1.2 (**Fig. S2a**). The cell cycle library exhibited a lower fitness distribution, consistent with cell cycle targets being enriched for essential genes. We then benchmarked the single-gene targeting performance of our genome-wide library against the *S. aureus* Lisbon CRISPRi Mutant Library, a collection of arrayed single-CRISPRi constructs targeting essential genes^26^, and the set of genes within the Nebraska Transposon

Mutant Library^30^ for which no transposon insertions were recovered. In both cases, the overlapping sets of genes targeted by our genome-wide library were highly enriched for low fitness (**Fig. S2b**).

We recovered 494,956 distinct spacer-pairs that allowed us to quantify 115,497 gene-level interactions. Overall, we observed a distribution of genetic interactions (*ε_ixj_*) that centered around 0 with a slight skew towards positive interactions (**Fig. 3a**). We saw high concordance between replicates, especially for all *ε_ixj_* with an absolute value > 1 (r = 0.93, **Fig. 3b, Fig. S2c**). Applying this *ε_ixj_* effect size and after setting the false-discovery rate to 0.05, we identified 874 positive and 97 negative, for a total of 971 genetic interactions (**Fig. 3c**).

**Fig. 3.**
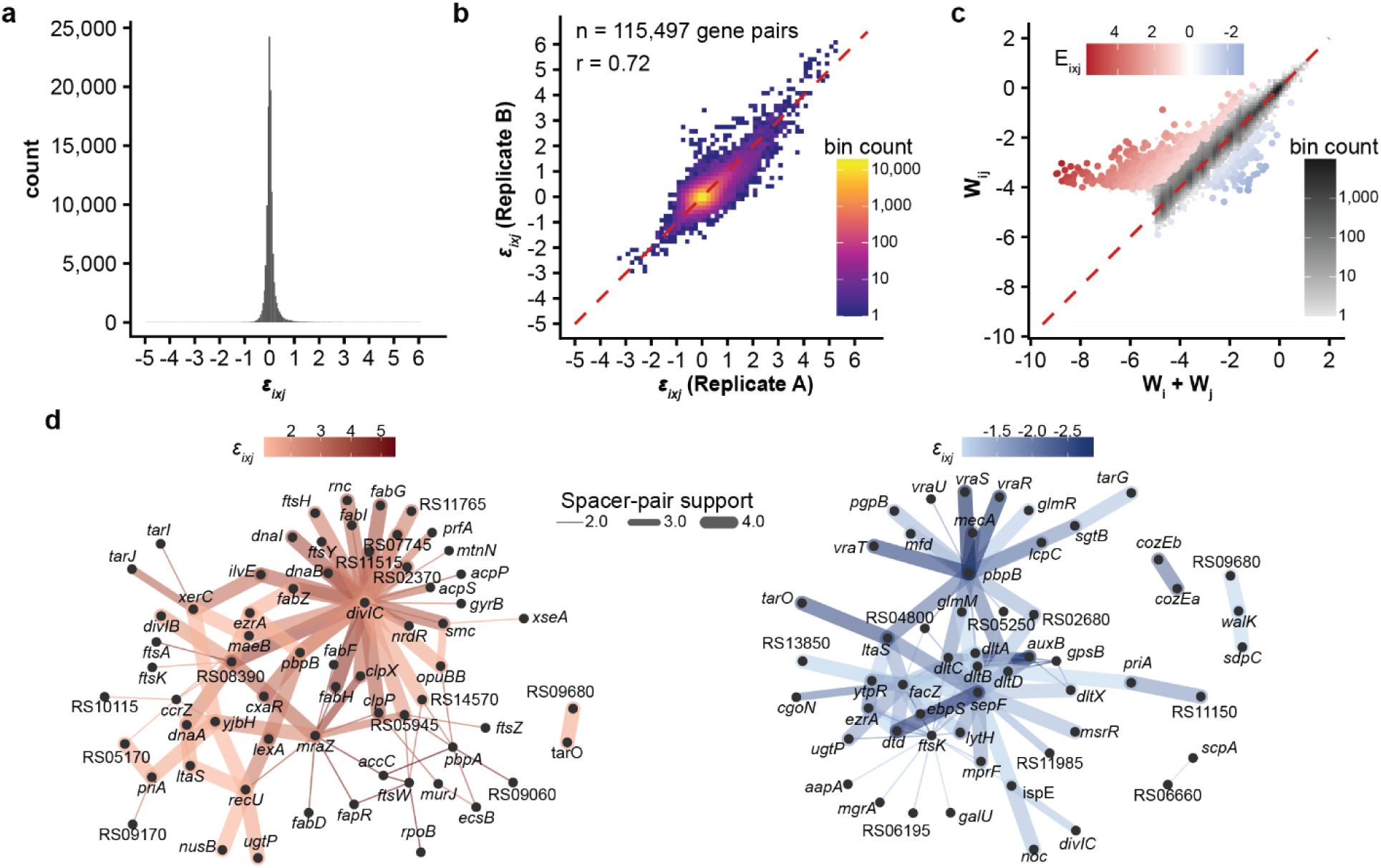
Global view of cell cycle genetic interactions. **a.** Histogram of genetic interaction scores (mean, n = 2 biological replicates). **b.** Concordance plot of interaction scores across two independent replicates (r is the Pearson correlation coefficient). **c.** Fitness plot of gene pairs comparing the expected additive fitness (W_i_ + W_j_) vs the observed fitness (W_ij_). Gene pairs with fdr < 0.05 are colored based on positive (red) and negative (blue) genetic interaction (*ε_ixj_*) values. Gene pairs with fdr ≥ 0.05 were binned and the pair counts are in greyscale based on the number of pairs within a bin. **d.** Network graphs of positive suppressive subtype (left) and negative (right) interactions. For clarity, only interactions supported by at least two spacer-pairs are shown.

### Positive interactions reveal a connection between fatty acid synthesis and cell division

The preponderance of positive interactions may in part be due to an over-representation of essential genes in the cell-cycle focused library (**Fig. S2a**), for which negative interactions are not as readily detectable. However, CRISPRi fitness defects also manifest over time as gene products are titrated below protein-specific thresholds needed to slow growth. Because relative abundance-based fitness integrates these time-dependent effects, unequal depletion kinetics may cause dual-gene silencing to resemble the more severe single gene knockdown. This can lead to spurious co-equal (W_ij_ ≈ W_i_ ≈ W_j_) or masking (W_ij_ ≈ min(W_i_, W_j_)) positive subtype interactions (**Fig. S3a,b**). We thereby restricted further analysis to suppressive (W_ij_ > min(W_i_, W_j_)) subtypes, which is equivalent to applying the Min neutrality model for quantifying genetic interactions^29^. This yielded 127 gene-pair interactions. (**Fig. 3d**, **Fig. S3c**).

Among these, genes for fatty acid and phospholipid biosynthesis, *accA, accC, accD, acpP, acpS*, *fabD*, *fabF, fabG*, *fabH, fabI, fabZ,* and *plsX*, were over-represented (**Fig. 4a, Fig. S4a**), encompassing 49 spacer pairs, and partially suppressed the deleterious effects of silencing a cluster of genes (**Fig. 4b**) that we refer to as FAPPI (fatty <u>a</u>cid and <u>p</u>hospholipid biosynthesis <u>p</u>ositive interactors). The FAPPI genes are involved in cell division and septal peptidoglycan synthesis—*ftsZ*, *ftsA*, *ezrA*, *ftsW*, *pbpA, divIC,* and *ftsL*—and include the transcriptional regulators *walR* and *mraZ* that respond to cell wall stress. We validated the interactions between *accC* with *ftsW* and *pbpA* (**Fig. 4c,d**) by monitoring the growth of homogenous populations encoding clonal sgRNA pairs (**Fig. 4e,f**).

**Fig. 4.**
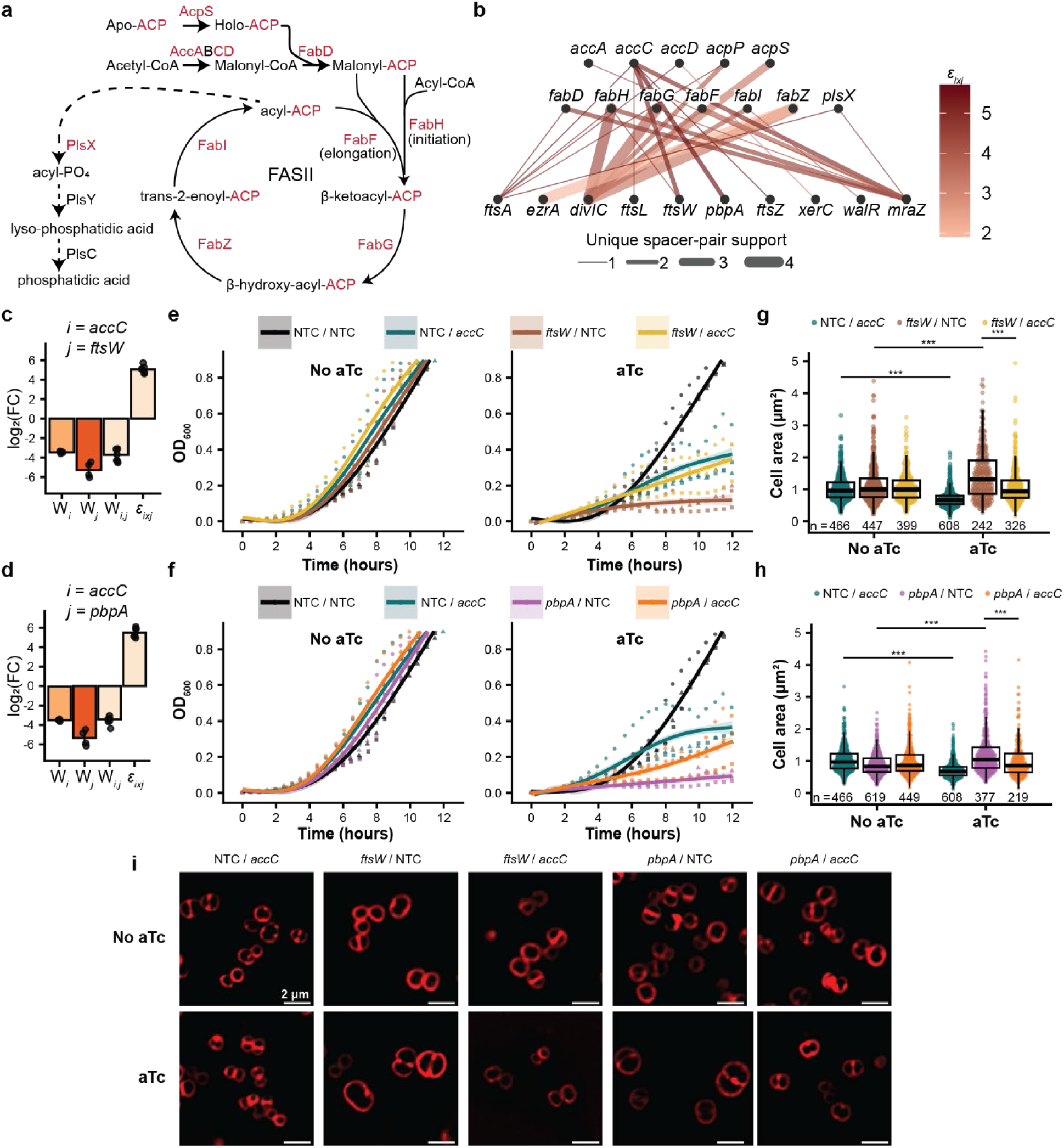
Silencing membrane biosynthesis genes suppress the fitness and cell enlargement defects of silencing FAPPI cell division genes. **a.** Fatty acid and phospholipid biosynthesis pathway in *S. aureus*. Genes for gene products in red exhibit suppressive interactions **b.** Network graph of positive genetic interactions (*ε_ixj_*) between fatty acid and phospholipid biosynthesis genes (top and middle rows) and the FAPPI set of cell division genes (bottom row). **c,d.** Bar graphs showing the mean fitness (W_i_, W_j_, W_ij_) and interactions (*ε_ixj_*) for all spacers targeting *accC* and (**c**) *ftsW* or (**d**) *pbpA.* e,f. Growth kinetics of clonal cells with sgRNA pairs targeting *accC* and (**e**) *ftsW* or (**f**) *pbpA* (n = 3 biological replicates; points show measurements and are shaped by replicate: circle, triangle, and square. GAM fitted curves, bands represent 95% confidence intervals). **g,h.** Box and violin plots of cell size for cells with sgRNA pairs targeting *accC* and (**g**) *ftsW* or (**h**) *pbpA* (***, p < 0.001, Wilcoxon signed-rank test). **i.** Representative microscopy images of cells stained with the membrane dye FM4-64 (scale bar: 2 µm; NTC: non-targeting control).

Our results suggested that decreasing flux through fatty acid and phospholipid biosynthesis can ameliorate the loss of cell cycle functions, which is analogous to what occurs in *Bacillus subtilis*^40^. Moreover in *B. subtilis*, cell size is controlled by fatty acid synthesis by limiting membrane capacity^41^. Intriguingly, disrupting genes within the FAPPI set commonly leads to large-cell phenotypes^42–49^. We therefore hypothesized that enlargement of cells, at least in part, is responsible for the reduced fitness of silencing FAPPI genes and that simultaneous blocking of fatty acids or phospholipid biosynthesis suppresses the fitness defect of FAPPI silencing by restricting cell enlargement. After imaging and quantifying the size of cells with clonal sgRNA pairs (**Fig. 4g-i**), we replicated the large cell phenotypes by silencing *ftsW* or *pbpA* alone. Conversely, silencing *accC* alone reduced cell size and, consistent with our hypothesis, dual silencing of *accC* with *ftsW* or *pbpA* led to a cell size distribution similar to control cells in which silencing was not induced.

### D-alanylation synthetic lethality with cell division genes

Among the negative interactions (**Fig. 3d, Fig. S4b**), we again detected an interaction between *tarO* and *ltaS* as well as the known interaction between the paralogs *cozEa* and *cozEb*^22^. In addition, *pbpB*, which encodes for the sole class A penicillin-binding protein (PBP) PBP2 in *S. aureus* and is the target of methicillin, was a hub of 20 interactions. These include *mecA*, which confers methicillin resistance by encoding the alternative PBP, PBP2a; *vraTRS* which encode for a three-component signaling system that detects PBP2 inhibition^50^ by β-lactams and vancomycin^51,52^; *glmR* and *glmM*, which biosynthesize the uridine diphosphate (UDP) carrier sugar in the early steps of peptidoglycan synthesis^53^; and *pptA*, which is homologous to *E. coli* phosphatase *pgpB* that is involved in recycling the lipid carrier undecaprenyl-diphoshpate^54,55^. Also notable, we detected interactions of *pbpB* with the CDSs BH7H15_RS05250 and B7H15_RS02680. The latter encodes for a member of the MOP exporter superfamily and has homology to *Bacillus subtilis* SpoVB^56^.

Unexpectedly, *pbpB* showed additional interactions with the *dlt* operon, *dltXABCD*, which encodes for a multi-protein pathway that modifies teichoic acids by attaching D-alanine esters^57–60^, and with the CDS (B7H15_RS04800) 48 base pairs upstream of *dltX* (n = 18 spacer pairs, **Fig 5a,b, Fig S5a,b**). Likewise, we detected robust interactions across the *dlt* operon with the divisome genes *gpsB*, *auxB*, *facZ*, *ftsK,* and *sepF* (n = 9, 18, 18, 9, and 18 spacer pairs, respectively, **Fig. 5b, Fig. S5c-g**), hereafter referred to as the DANI (<u>D</u>-<u>a</u>lanylation pathway <u>n</u>egative interactors) gene cluster. GpsB, which bundles FtsZ to aid in Z-ring constriction^61^, forms protein-protein interactions with the other DANI gene products PBP2^62,63^, FacZ^64^, and AuxB^65^, consistent with the notion that DANI genes function as a unit. However, *dlt* has not been functionally linked to *S. aureus* cell division. We therefore validated these interactions by monitoring growth of homogenous populations encoding clonal sgRNA pairs and, independently, silencing *dltA* in strains carrying transposon insertions that disrupt *gpsB* or *auxB* (**Fig 5c-3e**). Taken together, these findings identify a previously unrecognized functional connection between D-alanylation of teichoic acids and a discrete cell division module.

**Fig. 5.**
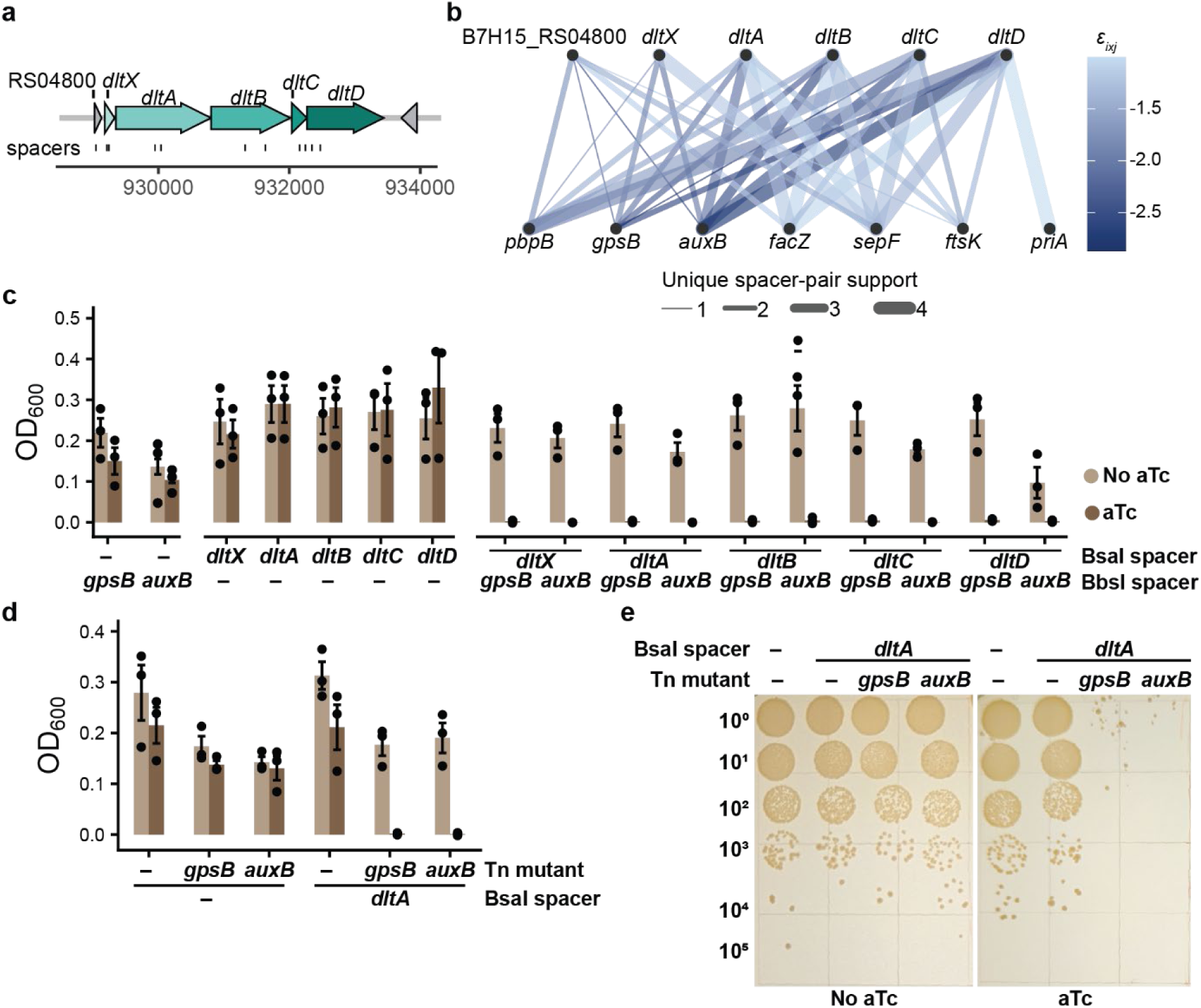
Interaction between genes responsible for D-alanylation of teichoic acids and the divisome components *pbpB*, *gpsB, facZ,* and *auxB*. **a**. Diagrams of the *dlt* operon structure and spacer targets (left). **b.** Network graph of cell cycle genes showing negative genetic interactions (*ε_ixj_*) with silencing *dlt* genes. **c.** Validation of genetic interactions by growth of selected *dlt* and *gpsB* or *auxB* clonal sgRNA pairs (n = 3 biological replicates; bars show mean ± standard error; for samples with aTc, p < 0.001 using ANOVA followed by Tukey’s HSD test comparing dual targeting to each single-targeting group). **d.** Validation of genetic interactions by growth using transposon insertion mutants of *gpsB* and *auxB* with a spacer targeting *dltA* (n = 3 biological replicates; bars show mean ± standard error; for samples with aTc, p < 0.05 for the interaction term by two-way ANOVA comparing wildtype vs transposon mutant alleles and *dltA* silencing). **e.** Ten-fold serial dilution and spot titers of *gpsB* and *auxB* transposon insertion mutants with a spacer targeting *dltA* on TSA plates supplemented with or without aTc.

## Discussion

We successfully developed a dual-CRISPRi platform for systematically identifying genetic interactions in MRSA, achieving genome-wide coverage for approximately 50 cell cycle genes of interest while deploying two independent spacers per target. Our validation efforts, confirming planned and unplanned known genetic interactions, establish this platform as a scalable and accessible approach for studying diverse pathways under a range of selective conditions.

Focusing our platform on characterized cell cycle genes, we identified hundreds of genetic interactions. Network analysis revealed two major clusters, one positive and one negative, that linked a distinct set of cell division genes with two bacterial cell surface pathways. The first, FAPPI, exhibited positive genetic interactions with genes involved in fatty acid and phospholipid biosynthesis and comprised a defined set of genes encoding divisome proteins. Many of these proteins are known to interact at the protein or functional level. Among these are the tubulin homolog FtsZ, which forms the Z-ring that constricts during cell division^66^, and its membrane anchor FtsA; as well as EzrA, which interacts with FtsZ and contributes to proper Z-ring assembly at the midcell^42,43^. The Z-ring serves as a scaffold for recruiting additional FAPPI gene products, such as the SEDS-bPBP pair FtsW-PBP1 and interacting proteins DivIC and FtsL ^42,43,46,67,68^, which altogether are responsible for septal peptidoglycan synthesis. In addition, this set included genes for the transcriptional regulators of cell division proteins MraZ, which controls the expression of *ftsL* in *B. subtilis*^69^, and the response regulator, WalR, of the two-component WalKR system that detects cell wall stress^70^.

Prior studies of individual FAPPI genes commonly reported enlarged cell phenotypes^42–49^. Our findings provide a mechanistic explanation in which membrane synthesis becomes excessive when FAPPI function is disrupted. We recapitulated the enlarged cell phenotype upon silencing *ftsW* and *pbpA*, which also caused severe population growth defects. Consistent with this model, restraining fatty acid biosynthesis by simultaneously silencing *accC* suppressed both phenotypes. Thus, at least in part, membrane overproduction underlies the negative effects of blocking cell division and indicates that *S. aureus*, similar to *B. subtilis*^40^, lacks sufficient regulatory capacity for relaying cell division defects to membrane biosynthetic pathways.

The second set, DANI, showed synthetic lethal interactions with silencing *dltXABCD* expression. This set also contained cell division genes, but with no direct overlap to the FAPPI set, encoding for PBP2, GpsB, FacZ, AuxB, SepF, and FtsK. As for the FAPPI set, multiple findings are consistent with a model in which DANI proteins work as a functional unit. GpsB localizes to the divisome where it bundles FtsZ but also serves as scaffolding protein forming protein-protein interactions with PBP2^62^, FacZ^64^, and AuxB^65^. In addition, SepF has been functionally implicated with FacZ because transposon insertions into either *sepF* or *facZ* were enriched in the same screen for membrane permeability and cell clumping^64^. Our results are also consistent with the sensitization to β-lactam antibiotics by the inhibition or genetic ablation of *dltA*^71,72^ for which the β-lactam target, PBP2, is a member of the DANI set.

The *dlt* operon is widely present in Firmicutes^73^ and encodes for DltXABCD that constitute a pathway for attaching D-alanine esters to LTA^74^, which can then be transferred to WTA. D-alanylation of teichoic acids reduces the overall negative surface charge that increases resistance to cationic antimicrobials^74,75^ and impacts metal ion homeostasis^59^. In addition, teichoic acids regulate the localization of cell wall hydrolases, although to our knowledge control of hydrolases by D-alanylation has not been reported. Because we added no cationic antimicrobials, we propose that either perturbation of metal ion homeostasis or dysregulated hydrolase activity may exacerbate the loss of DANI functions, thereby providing a mechanistic hypothesis that underlies the DANI-*dlt* synthetic lethal interaction.

In summary, our work establishes dual-CRISPRi as a scalable approach for genetic interaction profiling in *S. aureus*, uncovers two connections between cell division and the cell envelope pathways, provides a dataset for generating additional hypotheses of *S. aureus* cell cycle mechanisms, and serves as a roadmap for conducting genetic interaction profiling for poorly transformable bacterial pathogens.

## Methods

### Bacterial strains and growth conditions

*S. aureus* and *Escherichia coli* DC10B^76^ cells were routinely grown at 37°C with aeration in liquid tryptic soy broth (TSB; BD Bacto, #211822), liquid Brain Heart Infusion (BD Difco, #237500) media supplemented with 0.5 M sorbitol^77^ (BHIS), or on solid tryptic soy agar (TSA, TSB supplemented with 1.5% BD Difco granulated agar, #214510), supplemented, when necessary, with 20 µg/mL chloramphenicol (Cm), 10 µg/mL erythromycin (Erm), and 100 ng/mL of the inducer anhydrotetracycline (aTc). Strains, plasmids, and oligonucleotides used in this study are described in **Tables S3, S4, and S5**.

### Construction of S. aureus JE2 hsdR::dcas9-tetR

*S. aureus* JE2 *hsdR*::*dcas9-tetR* was engineered by allelic exchange^76^. Downstream and upstream *hsdR* homology arms were serially inserted into pIMAY by PCR and blunt-end ligations, after which the *Streptococcus pyogenes dcas9* was inserted under control of a *tetR*-P_tet_ promoter with 2x tetO operator cassette. The minus 10 sequences for P_tet_ and the *tetR* promoter were changed 5’-CATAAT-3’^31^ and 5’-TATAAT-3’^32^, yielding the allelic exchange vector pIMAY *hsdR*::*dcas9-tetR*. The plasmid sequence was confirmed by nanopore sequencing.

This vector was introduced into *S. aureus* JE2 by electroporation and allelic exchange was performed as before^78^. Cells from a single colony of JE2 *hsdR*::*dcas9-tetR* were confirmed by PCR amplification and nanopore sequencing of the integrated cassette, and by whole-genome sequencing using short-read sequencing.

### Transduction of transposon insertions

Transposon insertion mutants from the NTML library^30^ were transduced to JE2 *hsdR*::*dcas9-tetR* using bacteriophage ɸ11. Phage lysates of NTML transposon mutants were prepared using TSA supplemented with Erm10 and 5 mM CaCl_2_ and were then added to JE2 *hsdR*::*dcas9-tetR* cells. The sample was plated on TSA supplemented with Erm10 and 5 mM sodium citrate. The transposon insertion locus was confirmed by PCR and the *hsdR*::*dcas9-tetR* locus was confirmed by PCR and sequencing.

### Constructing the pSD14 dual-sgRNA vector

The pSD14 vector was constructed using elements from pIMAY (P_help_-*cat* for selection using Cm), pRB473 (for replication in *E. coli* or *S. aureus*), and a synthetic DNA fragment encompassing the dual sgRNA assembly sites. The P_help_-*cat* sequence was moved to between the two sgRNA sites to reduce recombination within cells between the identical sgRNA scaffold sequences. In addition, a BsmBI assembly site was introduced to allow for barcoding the BbsI sgRNA. The plasmid sequence was confirmed by nanopore sequencing.

### MRSA genome-wide spacer library design

All 20 bp spacers with a corresponding protospacer and adjacent PAM sequence within protein coding sequences (CDSs) were identified using the JE2 reference genome sequence and annotation (NZ_CP020619.1). These were filtered to remove spacers that 1) targeted the template strand, 2) had off-targets predicted by the CRISPRseek package^79^, 3) targeted overlapping CDSs, and 4) had a BsaI restriction site. For each CDS, up to two spacers were chosen among the remaining set with preference for spacers targeting the 5’ half of the CDS and by predicted efficacy score that was generated using the CRISPRseek package.

Non-targeting controls spacers were generated using random 20 bp sequences with ATCG probabilities of 0.35, 0.35, 0.15, and 0.15, respectively. Random spacers with up to 4 mismatches with the JE2 chromosome or with a BsaI restriction site were removed. Genome-wide and non-targeting control spacers were flanked with pSD14-specific overhangs, BsaI restriction sites, and 25 bp priming sites.

### Construction of pooled libraries with barcoded-sgRNAs at pSD14 BbsI site

Barcodes were randomly generated 15 nucleotides and filtered for distinct sequences with a Hamming distance of 3, sequences with homopolymers of length less than 3, GC content between 30 and 80%, and a predicted melting temperature between 30 and 55 °C. Oligonucleotides representing both strands of each barcode and spacer were synthesized to have overhangs after annealing that are compatible with pSD14 digested with BsmBI (NEB, #R0739) or BbsI (NEB, #R3733), respectively. Annealing was conducted by mixing 1 µM of each oligonucleotide strand in annealing buffer (10 mM Tris, 50 mM NaCl, 1 mM EDTA, pH 7.5), heating to 95 °C for 5 min, and allowing the mixture to gradually cool to 25°C over 20 mins. The annealed oligonucleotides were treated with T4 polynucleotide kinase (NEB, #M0201) and stored at -20°C. Annealed barcodes and spacers were assembled into pSD14 serially using Golden Gate assembly. The reactions consisted of 50 fmoles pSD14, 0.5 pmoles (1:10 vector to insert molar ratio) barcodes or spacers, and were incubated for four cycles of 37 °C (BbsI) or 42 °C (BsmBI) for 10 minutes followed by 16 °C for 5 minutes. The assemblies were then incubated at 65 °C for 20 minutes and 37 °C (BbsI) or 42 °C (BsmBI) for 10 minutes. The assembled plasmid was transformed into chemically competent *E. coli* DC10B, plasmid was isolated from a single colony and confirmed by restriction digest screening using BsmBI or BbsI, respectively. After serial barcode-spacer assemblies, confirmed plasmids were pooled in equal amounts (200 ng each).

The barcode-spacer correspondence was confirmed by nanopore sequencing. The pooled plasmid library was linearized by digestion with Eco53kI (NEB, #R0116) and sequenced by nanopore using the Premium PCR sequencing service provided by Plasmidsaurus. This service ligates sequencing adapters and yields full-length plasmid reads. The reads were filtered for lengths between 4,300 and 4,400 bp and for quality score > 20, resulting in 2,951 reads (∼26x coverage). Cutadapt^80^ was used to extract the 20 bp spacer sequence assembled at the BbsI site and the 15 bp barcode assembled at the BsmBI site, allowing for a sequencing error rate of 0.2. All barcodes matched with one distinct spacer sequence (>95% of reads) and any discrepancies between anticipated barcode/BbsI-spacer associations were corrected according to the sequencing results.

### Construction of pooled libraries with sgRNAs at pSD14 BsaI site

Pilot-scale and genome-wide spacer pooled oligonucleotides were synthesized by Twist Bioscience and amplified for 12 or 10 cycles, respectively, using Q5 High-Fidelity DNA Polymerase (NEB, #M0491), and the PCR products were purified using a silica spin column. The PCR product was assembled into the pSD14 by Golden Gate assembly with BsaI (NEB, #R3733) using 50 fmoles of vector in a 20 µL reaction volume at a 1:10 vector to insert molar ratio. To achieve sufficient coverage, a 200 µL reaction volume (500 fmoles pSD14) was used for assembling the genome-wide library. The assembly reactions were incubated for 30 cycles of 37 °C for 10 minutes and 16°C for 5 minutes. The assembly reaction was drop dialyzed and electroporated into *E. coli* DC10B cells (2.5 kV, 25 µF, 200 Ω, 0.1-cm gap cuvettes). Twelve parallel transformations were conducted for the genome-wide plasmid library. Immediately after electroporation, cells were recovered in BHIS at 37 °C with aeration for 1 hour, then added to 500 mL of TSB supplemented with Cm and cultured overnight at 37 °C with aeration. Plasmid DNA was isolated using a midiprep kit (ThermoFisher, #A35892). At 1 hour after electroporation, a sample was serially diluted and plated on TSA supplemented with Cm to determine the total number of transformants.

### Construction of clonal dual sgRNA pSD14 plasmids

Oligonucleotides for selected spacers were synthesized as above (Constructing pooled libraries of barcoded-sgRNAs at pSD14 BbsI site) to have compatible overhangs after annealing for the BbsI or BsaI sites of pSD14. Likewise, the assembly reactions were identical as described above. Assembled plasmids (**Table S6**) were used individually and not pooled.

### Dual sgRNA library transformation of *S. aureus* JE2 *hsdR*::*dcas9-tetR*

Electrocompetent JE2 *hsdR*::*dcas9-tetR* cells were prepared by growth in BHIS to an OD_600_ of approximately 0.600, washed three times with ultra-pure H_2_O, concentrated approximately 170-fold and resuspended with ultra-pure H_2_O. For genome-wide plasmid library, 35 µg of plasmid was drop-dialyzed against ultra-pure H_2_O and electroporated (2 kV, 25 µF, 100 Ω, 0.1-cm gap cuvettes, Gene Pulser Xcell, Bio-Rad Laboratories) into electrocompetent JE2 *hsdR*::*dcas9-tetR* cells via 7 parallel transformations (5 µg DNA/transformation). For each transformation, 1 mL of BHIS media was added to the cells and cultured at 37°C, 180 rpm for 1 hour. The total number of transformants was estimated 1 hour after electroporation by counting CFUs. The cultures were pooled and added to 500 mL of BHIS supplemented with Cm and grown for 6 hours at 37°C, 180 rpm. 400 mL of the culture was mixed with 600 mL fresh TSB supplemented with Cm, split into two parallel 500 mL cultures and supplemented with or without 100 ng/mL aTc and grown for 4 hours at 30°C, 180 rpm. Each culture was subsequently diluted 1:100 into fresh TSB supplemented with Cm ± aTc and grown for 16 hours at 30°C, 180 rpm. Cells were pelleted by centrifugation at 4,000 rpm for 30 minutes and stored at -80 °C. At every step, samples were plated on TSA and TSA supplemented with Cm to enumerate CFUs and check library coverage. For pilot scale library, 5 µg plasmid DNA and proportionally reduced media volumes (10-fold less) were used.

### Amplicon Sequencing

Plasmids were isolated after treating cells with lysostaphin using a plasmid Midi prep Kit (ThermoFisher, #A35892). The BsaI-spacer and barcode region was amplified from 200 ng of plasmid DNA from ± aTc samples using Q5 DNA polymerase for 12 cycles. For samples from the pilot-scale screen primers with partial Illumina adaptors and distinct indices for different conditions (± aTc) were used, and the amplicons were pooled and sequenced using the AmpX service from AmpSeq LLC, which provides >100,000 paired-end reads. For samples from the genome-wide screen, staggered (0-6 random nucleotides) primers with partial Illumina adaptors and distinct inline indicies for different conditions (± aTc) were used, and the amplicons were pooled and sequenced on a 2×75 Medium Output Flow Cell (75 GB) using an Element AVITI sequencer (AmpSeq LLC).

Reads were filtered for quality score ≥20 and the barcodes for the BbsI spacer and the BsaI spacer were extracted with an error rate set to 0.1. The extracted spacers were mapped to reference files, and the counts-per-million (CPM) was calculated for each spacer-pair before any downstream filtering. For single-target measurements, the CPM numerator was derived from the summed reads in which the other sgRNA position contained a non-targeting control spacer.

### Calculation of fitness and genetic interactions

Individual spacer pairs were filtered for counts ≥20 and ≥8 for -aTc and +aTc conditions, respectively, to reduce variance due to low-read counts. The fitness (W) of each strain was calculated by comparing its relative abundance in the presence and absence of inducer for each replicate

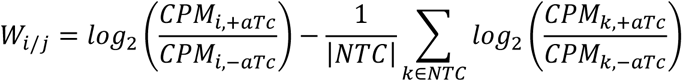

Where W_i_ or W_j_ is the fitness of targeting gene i or gene j by a spacer in the BsaI or BbsI position, respectively. The second half of the equation is the mean log_2_-fold-change among the set of non-targeting control spacers (NTC), which was used to adjust for compositional changes of the population upon induction with aTc due to loss of cells silencing essential genes.

The expected fitness of simultaneously silencing two targets were calculated using the Product model (additive after the log_2_ transformation):

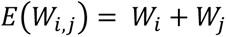

where W_i_ and W_j_ are the fitness values of single-gene targets quantified by the mean in which gene i or j is paired with a non-targeting control, and *E*(W_i,j_) is the expected fitness of silencing genes *i* and *j*. Genetic interactions were then derived by:

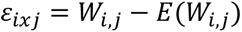

where W_ij_ is the observed fitness value of silencing genes *i* and *j*, and *ε_ixj_* is the genetic interaction between genes i and j.

We removed spacer pairs for which *E*(W_ij_) fell below a replicate-specific detection limit and W_ij_ was within 1 log_2_ unit of that detection limit. The latter allowed for the inclusion of spacer-pairs that were confidently recovered above the detection limit, regardless of if *E*(W_ij_) fell below it. The detection limit was set for each replicate to the 2.5th percentile of all single targeting fitness values (W_i_ and W_j_).

Positive interactions with a false discovery rate adjusted p-value < 0.05 were classified into subtypes using a tolerance of 0.5 log_2_ fitness units according to the following:

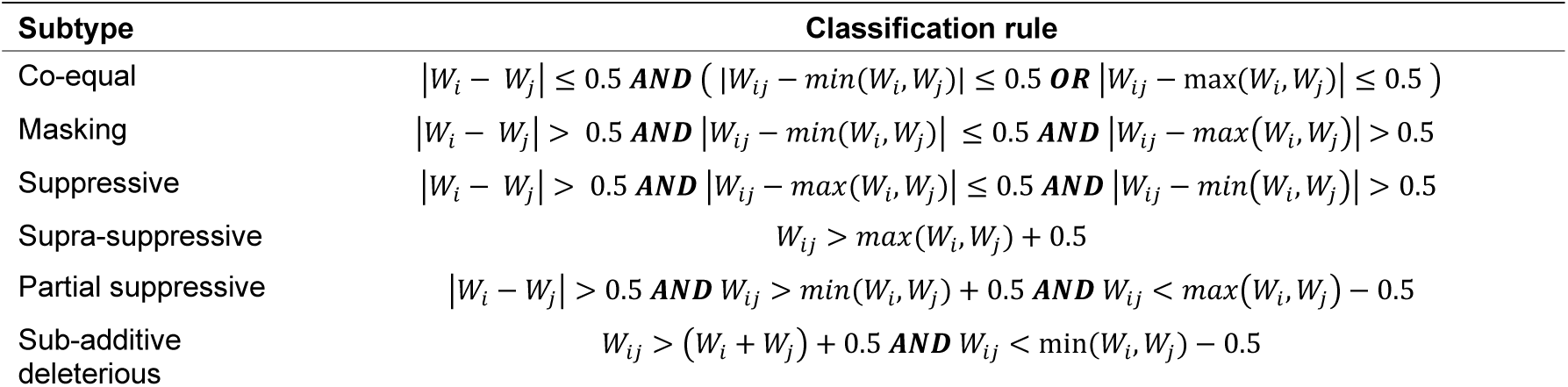

where max(W_i_, W_j_) and min(W_i_, W_j_) return the maximum or minimum value, respectively.

### Growth curves

Clonal JE2 *hsdR::dcas9-tetR* strains carrying distinct, individually assembled pSD14 constructs were grown overnight in TSB supplemented with Cm. Overnight cultures were diluted 1:500 into 2 mL fresh TSB supplemented with Cm and grown for 7 h at 37 °C with shaking. Culture densities were measured by OD_600_ in a 1-cm cuvette and the cultures were adjusted to OD_600_ = 1.2 as starting cell densities before a second 1:500 dilution into 2 mL TSB containing Cm, with or without aTc. Cultures were grown for 4 hours at 30 °C with shaking to induce CRISPRi. Following induction, cultures were diluted 1:100 into 200 µL fresh TSB containing Cm ± aTc in 96-well plates. OD_600_ was measured at the start of the assay and after 12 hours of growth at 30 °C with shaking, Experiments were performed on three independent days.

Clonal JE2 *hsdR::dcas9-tetR* strains carrying individually assembled pSD14 constructs targeting *accC, pbpA, ftsW* or the indicated gene pair were grown overnight in TSB supplemented with Cm. Culture densities were measured by OD_600_ in a 1-cm cuvette, and the cultures were diluted to an OD_600_ of 0.100 into 3 mL TSB supplemented with Cm and grown for 2 hours at 30 °C with shaking. The cultures were then diluted 16-fold into fresh TSB supplemented with Cm and divided evenly into two cultures. aTc was added to one culture and both cultures were transferred to a 96-well plate and incubated at 30 °C with shaking. Growth was monitored by OD_600_ in a plate reader (SpectraMax M2, Molecular Devices) at 30-minute intervals.

### Spot assays

Clonal JE2 *hsdR::dcas9-tetR* strains carrying the indicated single- or dual-targeting pSD14 constructs were grown overnight in TSB supplemented with Cm. Overnight cultures were diluted 1:10 into 2 mL fresh TSB supplemented with Cm and grown for 2 hours at 37 °C with shaking. Culture densities were measured by OD_600_ in a 1-cm cuvette and diluted to an OD_600_ of 1.2 before a second, 1:500 dilution into 2 mL TSB supplemented with Cm. Cultures were then serially 10-fold diluted, and 15 µL of each dilution was spotted onto TSA plates supplemented with Cm and with or without aTc. The plates were incubated at 30 °C for 36 hours.

### Microscopy

For fluorescence microscopy, clonal JE2 *hsdR::dcas9-tetR* strains carrying the indicated single- or dual-targeting pSD14 constructs were grown in TSB supplemented with Cm at 30°C with shaking at 250 rpm in a water bath. Overnight cultures were diluted into fresh TSB supplemented with Cm to an initial OD_600_ of 0.05 and grown to an OD_600_ of 0.2. Where indicated, expression was induced by addition of aTc, and cultures were incubated for an additional 6 hours at 30°C with shaking. Following incubation, 1 mL samples were harvested by centrifugation, washed with PBS, and resuspended in PBS containing 50 µg/mL FM4-64 (Invitrogen) to stain the cell membrane. Cells were mounted on 1% (w/v) agarose pads prepared in PBS and imaged using a DeltaVision Core microscope system (Applied Precision/GE Healthcare) equipped with a Photometrics CoolSnap HQ2 camera. The data were deconvolved using SoftWorx software as described previously^81^. For quantification of cell area, FM4-64 fluorescence images were segmented using Cellpose^82^ to generate individual cell regions of interest (ROIs). Segmentation masks were manually inspected and corrected when necessary. Cell area and other morphological parameters were subsequently measured from the resulting ROIs using Fiji.

### Statistics & Reproducibility

Statistical tests for genetic interactions of pooled libraries were performed on the gene level using the limma package for R to implement empirical Bayes^83^. The null hypothesis was set to 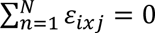, where n is an independent replicate and N is the total number of replicates. Multiple comparisons were controlled by setting the BH-FDR to 0.05. Graphs were created using the R language and R Studio IDE. The one-sided Wilcoxon rank sum test was used to test for differences in cell size. ANOVA was used to test for differences across groups and the Tukey Honestly Significant Difference method was used to test for pairwise differences.

## Acknowledgements

This work was funded by the University of Maryland Grand Challenges 2.0 Grants Program (to S.W.D.) and by University of Maryland startup funds (to S.W.D.) and the Intramural Research Program of the National Institutes of Health (NIH), National Cancer Institute, Center for Cancer Research (to K.S.R.). The contributions of the NIH authors are considered Works of the United States Government. The findings and conclusions presented in this paper are those of the authors and do not necessarily reflect the views of the NIH or the U.S. Department of Health and Human Services.

## Author Contributions Statement

**Sumana Bhowmick**: Conceptualization, Data Curation, Investigation, Validation, Visualization, Writing – original draft **Félix Ramos-León**: Data Curation, Formal Analysis, Investigation, Methodology, Writing – review & editing; **Kumaran S. Ramamurthi**: Writing – review & editing, Supervision; **Seth W. Dickey**: Conceptualization, Data Curation, Formal Analysis, Funding Acquisition, Investigation, Methodology, Project Administration, Supervision, Validation, Visualization, Writing – original draft, Writing – review & editing

## References

1. Pinho, M. G. & Foster, S. J. Cell Growth and Division of Staphylococcus aureus. Annu Rev Microbiol 78, 293–310 (2024).

2. Dewachter, L., Verstraeten, N., Fauvart, M. & Michiels, J. An integrative view of cell cycle control in Escherichia coli. FEMS Microbiol Rev 42, 116–136 (2018).

3. Meunier, A., Cornet, F. & Campos, M. Bacterial cell proliferation: from molecules to cells. FEMS Microbiol Rev 45, fuaa046 (2021).

4. Barbuti, M. D., Myrbråten, I. S., Morales Angeles, D. & Kjos, M. The cell cycle of Staphylococcus aureus: An updated review. Microbiologyopen 12, e1338 (2023).

5. Garcia, P. S. et al. A Comprehensive Evolutionary Scenario of Cell Division and Associated Processes in the Firmicutes. Mol Biol Evol 38, 2396–2412 (2021).

6. Ramos-León, F. & Ramamurthi, K. S. How do spherical bacteria regulate cell division? Biochem Soc Trans 53, 447–460 (2025).

7. Ramos-León, F. & Ramamurthi, K. S. Staphylococcus aureus as an emerging model to study bacterial cell division. J Biol Chem 301, 110343 (2025).

8. Pinho, M. G., Götz, F. & Peschel, A. Staphylococcus aureus: a model for bacterial cell biology and pathogenesis. J Bacteriol 207, e0010625 (2025).

9. Ramos-León, F. et al. PcdA promotes orthogonal division plane selection in Staphylococcus aureus. Nat Microbiol 9, 2997–3012 (2024).

10. Antimicrobial Resistance Collaborators. Global burden of bacterial antimicrobial resistance in 2019: a systematic analysis. Lancet 399, 629–655 (2022).

11. Turner, N. A. et al. Methicillin-resistant Staphylococcus aureus: an overview of basic and clinical research. Nat Rev Microbiol 17, 203–218 (2019).

12. Baryshnikova, A., Costanzo, M., Myers, C. L., Andrews, B. & Boone, C. Genetic interaction networks: toward an understanding of heritability. Annu Rev Genomics Hum Genet 14, 111–133 (2013).

13. Halder, V., McDonnell, B., Uthayakumar, D., Usher, J. & Shapiro, R. S. Genetic interaction analysis in microbial pathogens: unravelling networks of pathogenesis, antimicrobial susceptibility and host interactions. FEMS Microbiol Rev 45, fuaa055 (2021).

14. Typas, A. et al. High-throughput, quantitative analyses of genetic interactions in E. coli. Nat Methods 5, 781–787 (2008).

15. Dénéréaz, J. et al. Dual CRISPRi-seq for genome-wide genetic interaction studies identifies key genes involved in the pneumococcal cell cycle. Cell Syst 16, 101408 (2025).

16. Koo, B.-M. et al. Comprehensive genetic interaction analysis of the Bacillus subtilis envelope using double-CRISPRi. Cell Syst Koo, Byoung-Mo, Horia Todor, Jiawei Sun, et al. “Comprehensive Genetic Interaction Analysis of the Bacillus Subtilis Envelope Using Double-CRISPRi.” Cell Systems 16, no. 11 (2025): 101406. 10.1016/j.cels.2025.101406., 101406 (2025).

17. Zik, J. J. et al. Dual transposon sequencing profiles the genetic interaction landscape in bacteria. Science 389, eadt7685 (2025).

18. Oku, Y. et al. Pleiotropic roles of polyglycerolphosphate synthase of lipoteichoic acid in growth of Staphylococcus aureus cells. J Bacteriol 191, 141–151 (2009).

19. Swoboda, J. G., Campbell, J., Meredith, T. C. & Walker, S. Wall teichoic acid function, biosynthesis, and inhibition. Chembiochem 11, 35–45 (2010).

20. D’Elia, M. A. et al. Lesions in teichoic acid biosynthesis in Staphylococcus aureus lead to a lethal gain of function in the otherwise dispensable pathway. J Bacteriol 188, 4183–4189 (2006).

21. Joo, H.-S. et al. Mechanism of Gene Regulation by a Staphylococcus aureus Toxin. mBio 7, e01579–16 (2016).

22. Stamsås, G. A. et al. CozEa and CozEb play overlapping and essential roles in controlling cell division in Staphylococcus aureus. Mol Microbiol 109, 615–632 (2018).

23. Waldron, D. E. & Lindsay, J. A. Sau1: a novel lineage-specific type I restriction-modification system that blocks horizontal gene transfer into Staphylococcus aureus and between S. aureus isolates of different lineages. J Bacteriol 188, 5578–5585 (2006).

24. Monk, I. R. & Foster, T. J. Genetic manipulation of Staphylococci-breaking through the barrier. Front Cell Infect Microbiol 2, 49 (2012).

25. Feng, S. Y., Hauck, Y., Morgene, F., Mohammedi, R. & Mirouze, N. The complex regulation of competence in Staphylococcus aureus under microaerobic conditions. Commun Biol 6, 512 (2023).

26. Reed, P. et al. A CRISPRi-based genetic resource to study essential Staphylococcus aureus genes. mBio 15, e0277323 (2024).

27. Jiang, W., Oikonomou, P. & Tavazoie, S. Comprehensive Genome-wide Perturbations via CRISPR Adaptation Reveal Complex Genetics of Antibiotic Sensitivity. Cell 180, 1002–1017.e31 (2020).

28. Liu, X. et al. Genome-wide CRISPRi screens for high-throughput fitness quantification and identification of determinants for dalbavancin susceptibility in Staphylococcus aureus. mSystems 9, e0128923 (2024).

29. Mani, R., St Onge, R. P., Hartman, J. L., Giaever, G. & Roth, F. P. Defining genetic interaction. Proc Natl Acad Sci U S A 105, 3461–3466 (2008).

30. Fey, P. D. et al. A genetic resource for rapid and comprehensive phenotype screening of nonessential Staphylococcus aureus genes. mBio 4, e00537–12 (2013).

31. Helle, L. et al. Vectors for improved Tet repressor-dependent gradual gene induction or silencing in Staphylococcus aureus. Microbiology (Reading) 157, 3314–3323 (2011).

32. Corrigan, R. M. & Foster, T. J. An improved tetracycline-inducible expression vector for Staphylococcus aureus. Plasmid 61, 126–129 (2009).

33. Monk, I. R., Tree, J. J., Howden, B. P., Stinear, T. P. & Foster, T. J. Complete Bypass of Restriction Systems for Major Staphylococcus aureus Lineages. mBio 6, e00308–00315 (2015).

34. Markley, A. L., Begemann, M. B., Clarke, R. E., Gordon, G. C. & Pfleger, B. F. Synthetic biology toolbox for controlling gene expression in the cyanobacterium Synechococcus sp. strain PCC 7002. ACS Synth Biol 4, 595–603 (2015).

35. Ouyang, S. & Lee, C. Y. Transcriptional analysis of type 1 capsule genes in Staphylococcus aureus. Mol Microbiol 23, 473–482 (1997).

36. Santa Maria, J. P., et al. Compound-gene interaction mapping reveals distinct roles for Staphylococcus aureus teichoic acids. Proc Natl Acad Sci U S A 111, 12510–12515 (2014).

37. Kanampalliwar, A. et al. The role of lipoteichoic acid in *Staphylococcus aureus* cell wall integrity. bioRxiv 2025.01.16.633316 (2025) doi:10.1101/2025.01.16.633316.

38. Choe, D. et al. Genome-scale analysis of Methicillin-resistant Staphylococcus aureus USA300 reveals a tradeoff between pathogenesis and drug resistance. Sci Rep 8, 2215 (2018).

39. Prados, J., Linder, P. & Redder, P. TSS-EMOTE, a refined protocol for a more complete and less biased global mapping of transcription start sites in bacterial pathogens. BMC Genomics 17, 849 (2016).

40. Willdigg, J. R., Patel, Y. & Helmann, J. D. A Decrease in Fatty Acid Synthesis Rescues Cells with Limited Peptidoglycan Synthesis Capacity. mBio 14, e0047523 (2023).

41. Vadia, S. et al. Fatty Acid Availability Sets Cell Envelope Capacity and Dictates Microbial Cell Size. Curr Biol 27, 1757–1767.e5 (2017).

42. Steele, V. R., Bottomley, A. L., Garcia-Lara, J., Kasturiarachchi, J. & Foster, S. J. Multiple essential roles for EzrA in cell division of Staphylococcus aureus. Mol Microbiol 80, 542–555 (2011).

43. Jorge, A. M., Hoiczyk, E., Gomes, J. P. & Pinho, M. G. EzrA contributes to the regulation of cell size in Staphylococcus aureus. PLoS One 6, e27542 (2011).

44. Tinajero-Trejo, M. et al. The Staphylococcus aureus cell division protein, DivIC, interacts with the cell wall and controls its biosynthesis. Commun Biol 5, 1228 (2022).

45. Tinajero-Trejo, M. et al. Control of morphogenesis during the Staphylococcus aureus cell cycle. Sci Adv 11, eadr5011 (2025).

46. Reichmann, N. T. et al. SEDS-bPBP pairs direct lateral and septal peptidoglycan synthesis in Staphylococcus aureus. Nat Microbiol 4, 1368–1377 (2019).

47. Pinho, M. G. & Errington, J. Dispersed mode of Staphylococcus aureus cell wall synthesis in the absence of the division machinery. Mol Microbiol 50, 871–881 (2003).

48. Henriksen, C. et al. The ClpX chaperone and a hypermorphic FtsA variant with impaired self-interaction are mutually compensatory for coordinating Staphylococcus aureus cell division. Mol Microbiol 121, 98–115 (2024).

49. Wacnik, K. et al. Penicillin-Binding Protein 1 (PBP1) of Staphylococcus aureus Has Multiple Essential Functions in Cell Division. mBio 13, e0066922 (2022).

50. Fernandes, P. B., Reed, P., Monteiro, J. M. & Pinho, M. G. Revisiting the Role of VraTSR in Staphylococcus aureus Response to Cell Wall-Targeting Antibiotics. J Bacteriol 204, e0016222 (2022).

51. Yin, S., Daum, R. S. & Boyle-Vavra, S. VraSR two-component regulatory system and its role in induction of pbp2 and vraSR expression by cell wall antimicrobials in Staphylococcus aureus. Antimicrob Agents Chemother 50, 336–343 (2006).

52. Gardete, S., Wu, S. W., Gill, S. & Tomasz, A. Role of VraSR in antibiotic resistance and antibiotic-induced stress response in Staphylococcus aureus. Antimicrob Agents Chemother 50, 3424–3434 (2006).

53. Barreteau, H. et al. Cytoplasmic steps of peptidoglycan biosynthesis. FEMS Microbiol Rev 32, 168–207 (2008).

54. Kumar, S., Mollo, A., Kahne, D. & Ruiz, N. The Bacterial Cell Wall: From Lipid II Flipping to Polymerization. Chem Rev 122, 8884–8910 (2022).

55. Kawakami, N. & Fujisaki, S. Undecaprenyl phosphate metabolism in Gram-negative and Gram-positive bacteria. Biosci Biotechnol Biochem 82, 940–946 (2018).

56. Vasudevan, P., McElligott, J., Attkisson, C., Betteken, M. & Popham, D. L. Homologues of the Bacillus subtilis SpoVB protein are involved in cell wall metabolism. J Bacteriol 191, 6012–6019 (2009).

57. Schultz, B. J., Snow, E. D. & Walker, S. Mechanism of D-alanine transfer to teichoic acids shows how bacteria acylate cell envelope polymers. Nat Microbiol 8, 1318–1329 (2023).

58. Wood, B. M., Santa Maria, J. P., Matano, L. M., Vickery, C. R. & Walker, S. A partial reconstitution implicates DltD in catalyzing lipoteichoic acid d-alanylation. J Biol Chem 293, 17985–17996 (2018).

59. Neuhaus, F. C. & Baddiley, J. A continuum of anionic charge: structures and functions of D-alanyl-teichoic acids in gram-positive bacteria. Microbiol Mol Biol Rev 67, 686–723 (2003).

60. Koch, H. U., Döker, R. & Fischer, W. Maintenance of D-alanine ester substitution of lipoteichoic acid by reesterification in Staphylococcus aureus. J Bacteriol 164, 1211–1217 (1985).

61. Eswara, P. J. et al. An essential Staphylococcus aureus cell division protein directly regulates FtsZ dynamics. Elife 7, e38856 (2018).

62. Costa, S. F. et al. The role of GpsB in Staphylococcus aureus cell morphogenesis. mBio 15, e0323523 (2024).

63. Sutton, J. A. F. et al. The roles of GpsB and DivIVA in Staphylococcus aureus growth and division. Front Microbiol 14, 1241249 (2023).

64. Bartlett, T. M. et al. FacZ is a GpsB-interacting protein that prevents aberrant division-site placement in Staphylococcus aureus. Nat Microbiol 9, 801–813 (2024).

65. Sisley, T. A. et al. AuxB interacts directly with GpsB and PknB to coordinate cell envelope processes that contribute to intrinsic antibiotic resistance in Staphylococcus aureus. mBio 16, e0185825 (2025).

66. Barrows, J. M. & Goley, E. D. FtsZ dynamics in bacterial division: What, how, and why? Curr Opin Cell Biol 68, 163–172 (2021).

67. Käshammer, L. et al. Cryo-EM structure of the bacterial divisome core complex and antibiotic target FtsWIQBL. Nat Microbiol 8, 1149–1159 (2023).

68. Nguyen, H. T. V. et al. Structure of the heterotrimeric membrane protein complex FtsB-FtsL-FtsQ of the bacterial divisome. Nat Commun 14, 1903 (2023).

69. White, M. L., Hough-Neidig, A., Khan, S. J. & Eswara, P. J. MraZ Transcriptionally Controls the Critical Level of FtsL Required for Focusing Z-Rings and Kickstarting Septation in Bacillus subtilis. J Bacteriol 204, e0024322 (2022).

70. Sharkey, L. K. R. et al. The two-component system WalKR provides an essential link between cell wall homeostasis and DNA replication in Staphylococcus aureus. mBio 14, e0226223 (2023).

71. Coupri, D. et al. Inhibition of d-alanylation of teichoic acids overcomes resistance of methicillin-resistant Staphylococcus aureus. J Antimicrob Chemother 76, 2778–2786 (2021).

72. Brown, S. et al. Methicillin resistance in Staphylococcus aureus requires glycosylated wall teichoic acids. Proc Natl Acad Sci U S A 109, 18909–18914 (2012).

73. Jordan, S., Hutchings, M. I. & Mascher, T. Cell envelope stress response in Gram-positive bacteria. FEMS Microbiol Rev 32, 107–146 (2008).

74. Peschel, A. et al. Inactivation of the dlt operon in Staphylococcus aureus confers sensitivity to defensins, protegrins, and other antimicrobial peptides. J Biol Chem 274, 8405– 8410 (1999).

75. Peschel, A., Vuong, C., Otto, M. & Götz, F. The D-alanine residues of Staphylococcus aureus teichoic acids alter the susceptibility to vancomycin and the activity of autolytic enzymes. Antimicrob Agents Chemother 44, 2845–2847 (2000).

76. Monk, I. R., Shah, I. M., Xu, M., Tan, M. W. & Foster, T. J. Transforming the untransformable: application of direct transformation to manipulate genetically Staphylococcus aureus and Staphylococcus epidermidis. mBio 3, (2012).

77. Chen, Y. E. et al. Engineered skin bacteria induce antitumor T cell responses against melanoma. Science 380, 203–210 (2023).

78. Dickey, S. W., Burgin, D. J., Huang, S., Maguire, D. & Otto, M. Two transporters cooperate to secrete amphipathic peptides from the cytoplasmic and membranous milieus. Proc. Natl. Acad. Sci. U.S.A. 120, e2211689120 (2023).

79. Zhu, L. J., Holmes, B. R., Aronin, N. & Brodsky, M. H. CRISPRseek: a bioconductor package to identify target-specific guide RNAs for CRISPR-Cas9 genome-editing systems. PLoS One 9, e108424 (2014).

80. Kechin, A., Boyarskikh, U., Kel, A. & Filipenko, M. cutPrimers: A New Tool for Accurate Cutting of Primers from Reads of Targeted Next Generation Sequencing. J Comput Biol 24, 1138–1143 (2017).

81. Tan, I. S., Weiss, C. A., Popham, D. L. & Ramamurthi, K. S. A Quality-Control Mechanism Removes Unfit Cells from a Population of Sporulating Bacteria. Dev Cell 34, 682–693 (2015).

82. Pachitariu, M., Rariden, M. & Stringer, C. Cellpose-SAM: superhuman generalization for cellular segmentation. bioRxiv 2025.04.28.651001 (2025) doi:10.1101/2025.04.28.651001.

83. Ritchie, M. E. et al. limma powers differential expression analyses for RNA-sequencing and microarray studies. Nucleic Acids Res 43, e47 (2015).

