## Supplementary information for "A genome-wide genetic interaction platform for MRSA reveals connections between cell division and the cell envelope"

**for**

#### List of supplementary information

**Supplementary Figure 1: Spacer-level concordance of fitness ($W_i$, $W_j$, $W_{ij}$) and genetic interactions ($\varepsilon_{ixj}$) using the pilot-scale library.**

**Supplementary Figure 2: Global view of gene-level fitness and spacer-level concordance.**

**Supplemental Figure 3: Classifying positive interactions into subtypes.**

**Supplemental Figure 4 Genome-wide genetic interactions with the cell cycle.**

**Supplemental Figure 5: Genetic interactions between DANI genes and *dltXABCD*.**

**Supplementary Table 1. Population size and library coverage for the genome-wide screen.**

**Supplementary Table 2. Genome-wide library sequencing coverage.**

**Supplementary Table 3. Bacterial strains used in this study.**

**Supplementary Table 4. Plasmids used in this study.**

**Supplementary Table 5. Oligonucleotides used in this study.**

**Supplementary Table 6. pSD14 plasmids used for gene silencing in clonal populations.**

**Supplementary References**

### Supplementary Figures and Legends

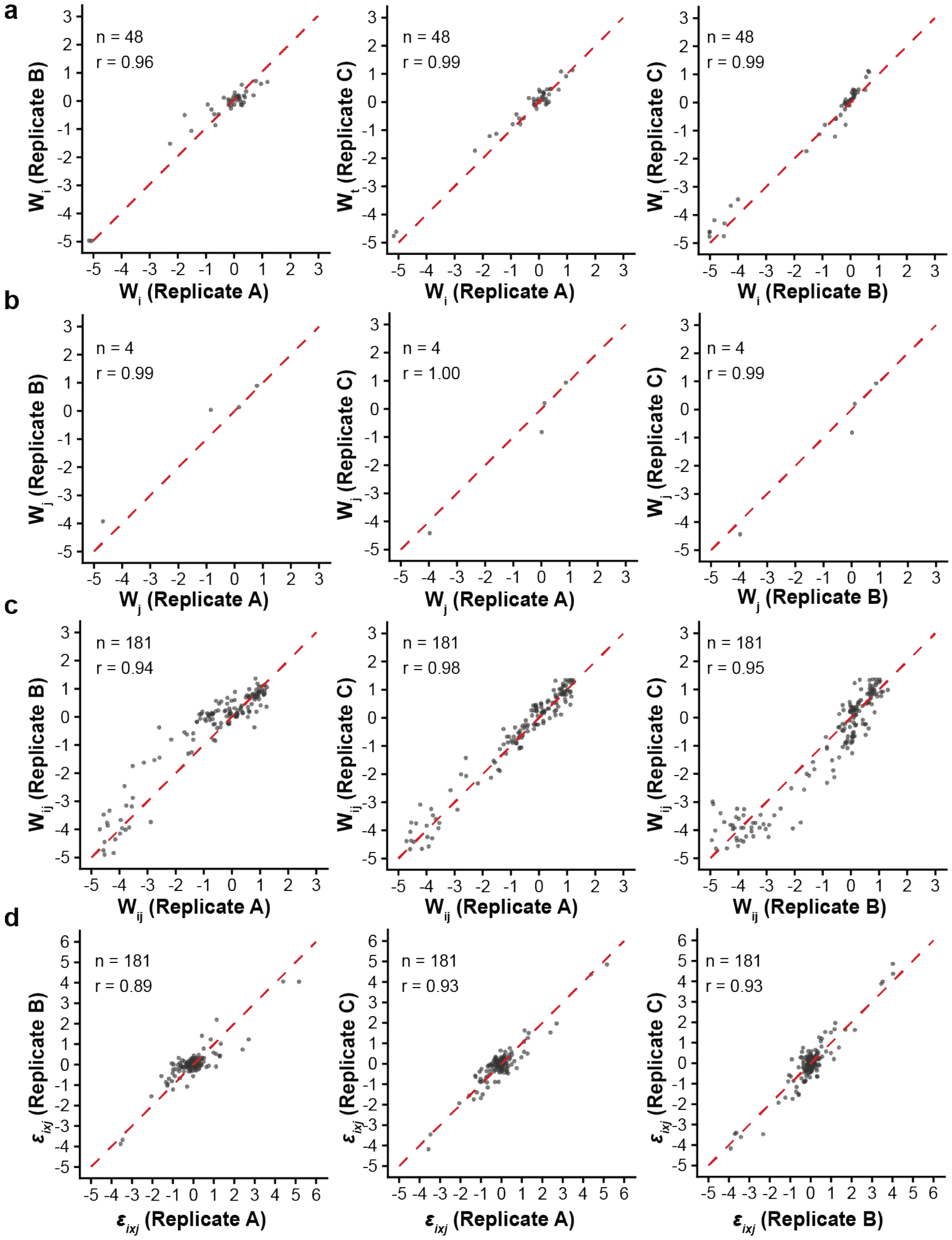


**Supplemental Figure 1: Spacer-level concordance of fitness ($W_i$, $W_j$, $W_{ij}$) and genetic interactions ($\varepsilon_{ixj}$) using the pilot-scale library. a,b.** $W_i$ (**a**) and $W_j$ (**b**) represent targeting spacers in the BsaI and BbsI positions, respectively. **c.** $W_{ij}$ represents targeting spacers in both the BsaI and BbsI positions. **d.** $\varepsilon_{ixj}$ represents the calculated interaction between targeting spacer pairs. n, number of distinct spacers or spacer pairs and r, the Pearson's correlation value between replicates.

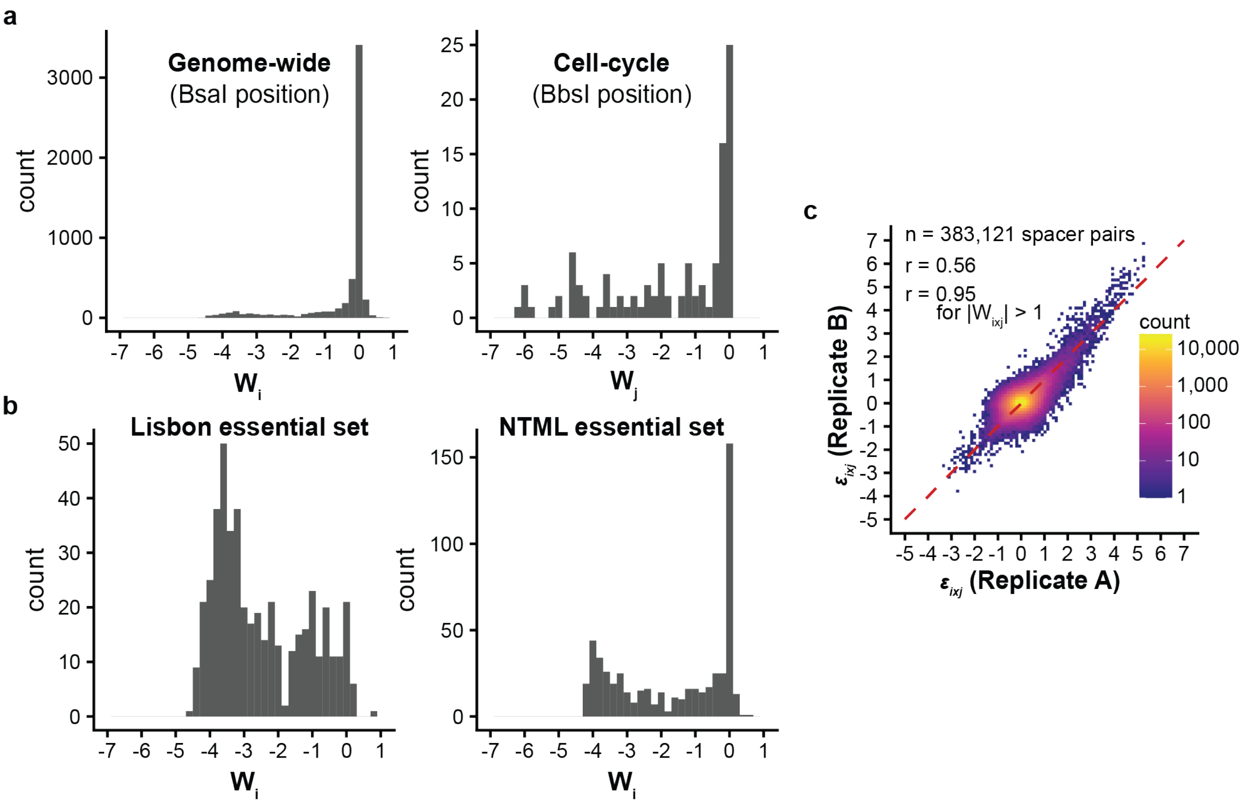


**Supplemental Figure 2: Global view of gene-level fitness and spacer-level concordance. a.** Single gene targeting fitness distribution for genome-wide targets (BsaI position, $W_i$) and cell cycle targets (BbsI position, $W_j$). Genes targeted by cell-cycle library were enriched for low fitness in comparison to the genome-wide library ($p < 0.0001$, Wilcoxon rank sum test) **b.** Single gene targeting fitness ($W_i$) for the set of genes targeted in the (left) Lisbon CRISPRi Mutant Library (1) or (right) the set without transposon insertions in the Nebraska Transposon Mutant Library (2). For both sets, genes targeted by the genome-wide library were enriched for low fitness ($p < 0.0001$, Wilcoxon rank sum test) **c.** Concordance plot of interaction scores at the spacer-pair level across two independent replicates**.**

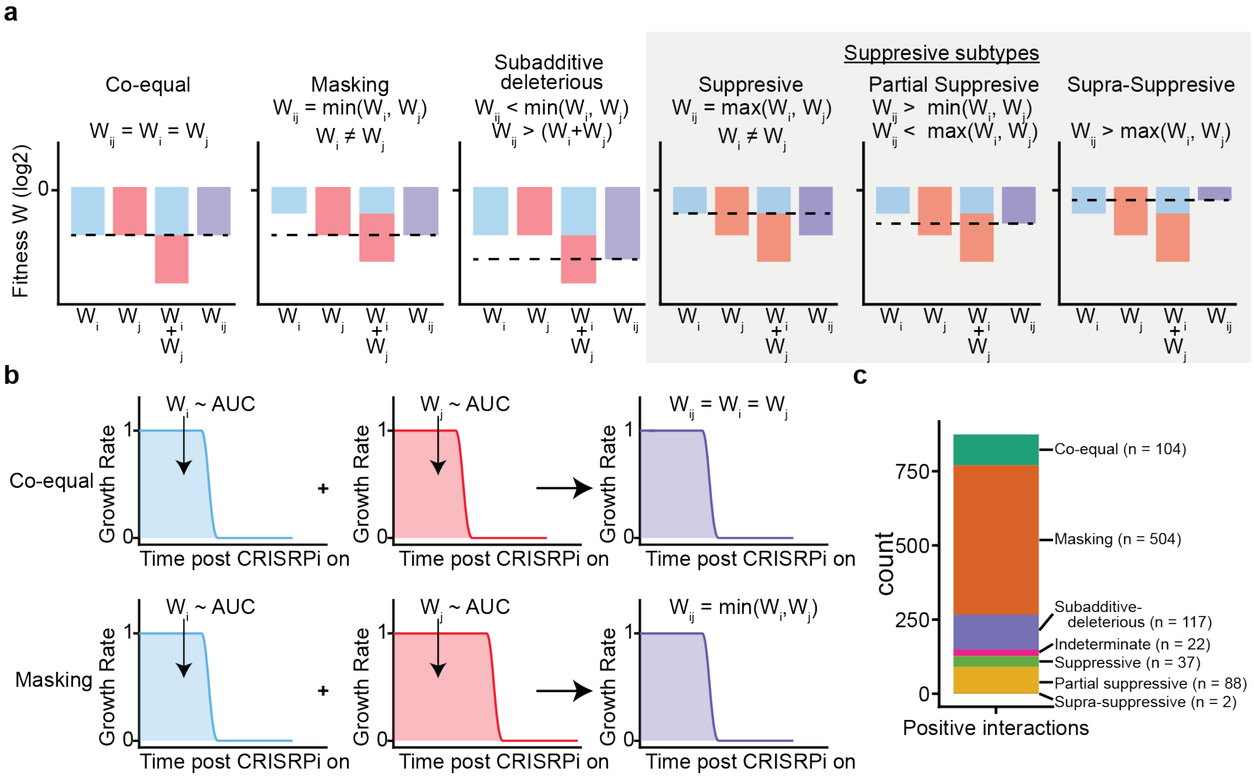


47 **Supplemental Figure 3: Classifying positive interactions into subtypes. a.** Positive genetic interaction subtypes based on comparisons between $W_i$, $W_j$, $W_i + W_j$ (the expected fitness of dual-silencing), and $W_{ij}$. **b.** Hypothetical cases in which silencing two distinct genes results in time-dependent growth arrest (relative growth rate = 0) after initiating CRISPRi silencing of gene expression, which results in spurious co-equal (top) or masking (bottom) positive subtype interactions without invoking a biologically relevant genetic interaction ($\varepsilon_{ixj}$). **c.** Stacked bar chart of positive subtype interactions (gene-level) using the cell cycle and genome wide dual silencing libraries.

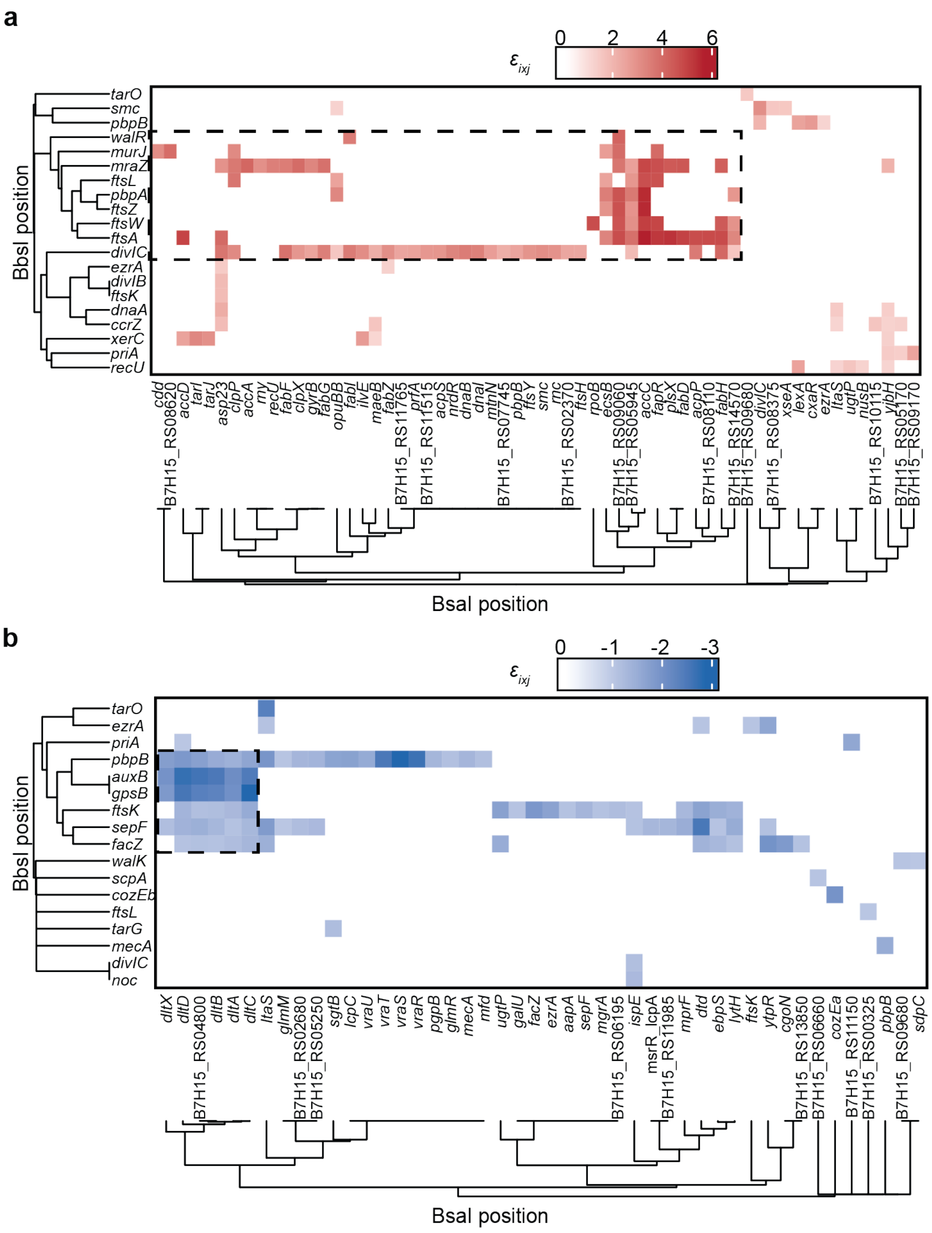


**Supplemental Figure 4 Genome-wide genetic interactions with the cell cycle. a,b.** Heatmap and clustering of (**a**) positive suppressive subtype interactions and (**b**) negative interactions. Targets in one position were clustered by Spearman rank correlation of genetic interactions ($\varepsilon_{ixj}$) with targets in the opposite position. All values where $|\varepsilon_{ixj}| < 1$ were set to 0. Dashed bounding boxes highlight clusters of interactions chosen for follow up analyses.

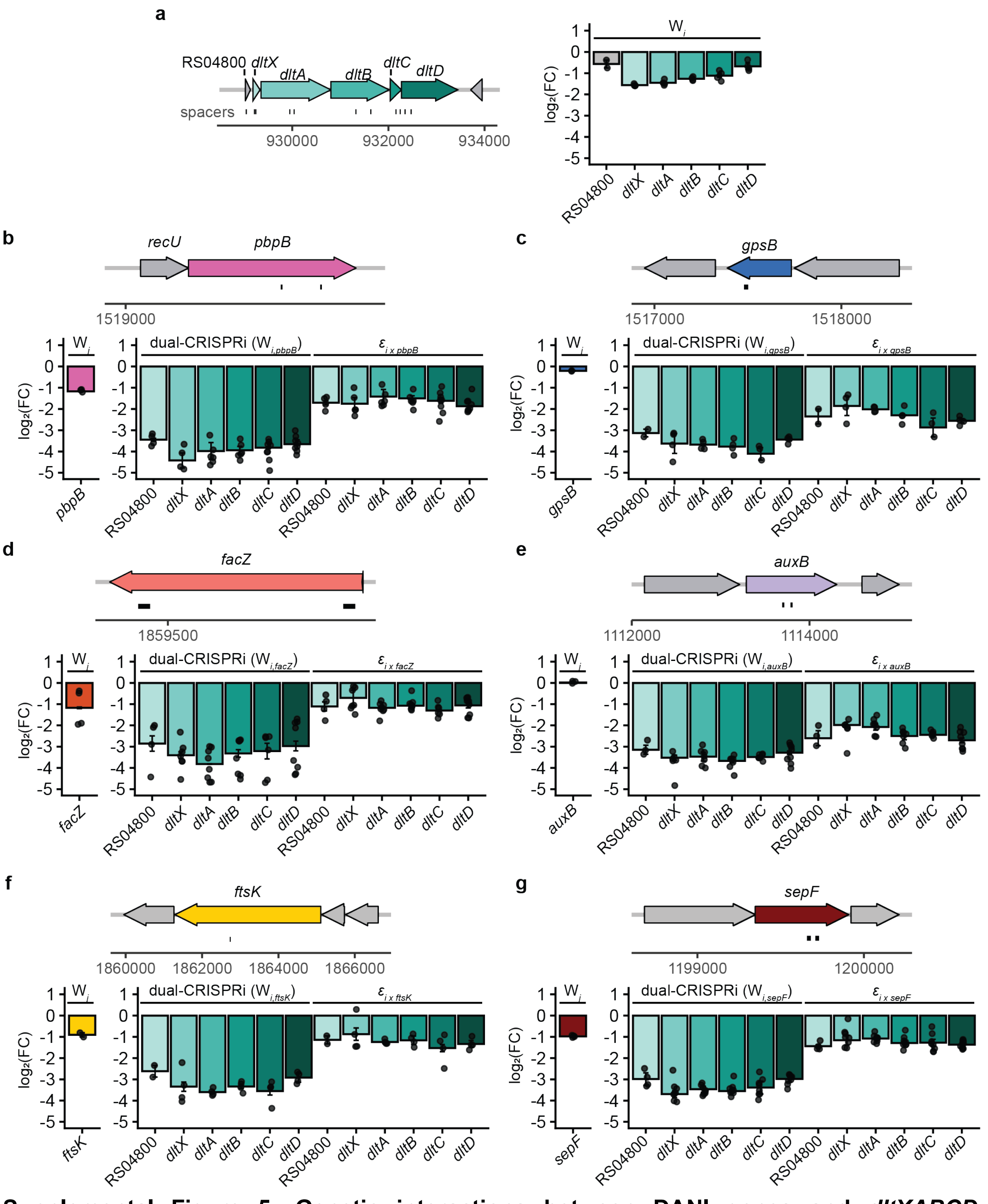


61 **Supplemental Figure 5: Genetic interactions between DANI genes and *dltXABCD*.**
62 Interaction between genes responsible for D-alanylation of teichoic acids and the divisome
63 components *pbpB*, *gpsB*, *facZ*, and *auxB*. **a**. Diagrams of the *dlt* operon structure and spacer

targets (left). Bar graphs of the fitness ($W_i$) targeting *dlt* genes. **b-g.** Diagrams of the (top) gene organization and (bottom) bar graphs of fitness targeting a single gene ($W_j$) or fitness and interactions targeting dual-genes $W_{ij}$) and interactions ($\varepsilon_{ixj}$) (right) for the (**b**) *pbpB*, (**c**) *gpsB*, (**d**) *facZ*, (**e**) *auxB*, (**f**), *ftsK*, and (**g**) *sepF*. Error bars summarize the mean and standard error across (n = 2) biological replicates.

### Supplementary Tables

#### Supplementary Table 1. Population size and library coverage for the genome-wide screen

| Bacterium | Replicate | Experimental Step | Total CFUs* | Library Coverage |
|---|---|---|---:|---:|
| *E. coli* | 1 and 2** | 1 hour after transformation | $1.1*10^{9}$ | 1,810x |
| *S. aureus* | 1 | 1 hour after transformation | $5.1*10^{*8}$ | 860x |
| *S. aureus* | 1 | 6 hours after pre-induction culture (BHIS) | $3.9*10^{9}$ | 6,470x |
| *S. aureus* | 1 | 4 hours after post-induction culture (without aTc) | $5.9*10^{9}$ | 9,930x |
| *S. aureus* | 1 | 4 hours after post-induction culture (with aTc) | $5.7*10^{9}$ | 9,500x |
| *S. aureus* | 2 | 1 hourr after transformation | $2.4*10^{8}$ | 410x |
| *S. aureus* | 2 | 6 hours after pre-induction culture (BHIS) | $6.6*10^{8}$ | 1,110x |
| *S. aureus* | 2 | 4 hours after post-induction culture (without aTc) | $1.2*10^{9}$ | 2,150x |
| *S. aureus* | 2 | 4 hours after post-induction culture (with aTc) | $4.8*10^{8}$ | 810x |

*Total CFUs was calculated from the CFUs/mL multiplied by the culture volume.

**Plasmid library isolated from *E. coli* served as the source for both replicates in *S. aureus*

#### Supplementary Table 2. Genome-wide library sequencing coverage

| Replicate | Condition | Total Reads | Sequencing coverage (Reads/Library diversity) |
|---|---|---|---|
| 1 | With aTc | 60,923,319 | 102 |
| 1 | Without aTc | 180,766,104 | 303 |
| 2 | With aTc | 196,321,939 | 330 |
| 2 | Without aTc | 225,680,474 | 379 |

**Supplementary Table 3. Bacterial strains used in this study.**

| Bacterial strain name | Description | Source |
|---|---|---|
| *Escherichia coli* DC10B | *dcm*[-] K12 strain used for subcloning and generating pSD14 plasmid libraries. | (3) |
| *Staphylococcus aureus* JE2 | USA300 MRSA strain derivative cured of native plasmids | (2) |
| *S. aureus* JE2 *hsdR::dcas9-tetR* | JE2 for inducible expression of *dcas9* | This study |
| *S. aureus* JE2 *gpsB*:Tn | NE1357 | (2) |
| *S. aureus* JE2 *hsdR::dcas9-tetR gpsB::*Tn | NE1357 derivative for inducible expression of *dcas9* | This study |
| *S. aureus* JE2 *auxB*:Tn | NE1420 | (2) |
| *S. aureus* JE2 *hsdR::dcas9-tetR auxB::*Tn | NE1420 derivative for inducible expression of *dcas9* | This study |

**Supplementary Table 4. Plasmids used in this study**

| Plasmid Name | Description/Purpose | Source |
|---|---|---|
| pIMAY | Staphylococcal allelic exchange vector; source of $P_{help}$-*cat* for pSD14; source of *tetR*-$P_{tet}$ for pIMAY *hsdR::dcas9-tetR* | (3) |
| pJMP1 | Source of *Streptococcus pyogenes dcas9* for pIMAY *hsdR::dcas9-tetR* | (4) |
| pIMAY *hsdR::dcas9-tetR* | Replace the *S. aureus hsdR* locus with the *tetR-dcas9* cassette | This study |
| pRB473 | Source of Gram-negative and Gram-positive plasmid replication for pSD14 | (5) |
| pSD14* | Dual sgRNA delivery vector | This study |

*See Supplementary Table S6 for list of assembled pSD14 plasmids and the corresponding spacer sequences.

#### Supplementary Table 5. Oligonucleotides used in this study.

| Oligonucleotide Name | Sequence (5'-3') | Purpose |
|---|---|---|
| 1kb_us_hsdR_forward | ATTTGCGACGCATGATG | Amplify *hsdR* upstream homology arm |
| delta_72bp_us_hsdR_reverse | ATCTTCAAACGTAAATTTATTCTAATTTTATTGTC | |
| delta_hsdR_stop_forward | TAATGATTCAGCCCCCTC | Amplify *hsdR* downstream homology arm |
| hsdR_ds_1kb_reverse | ATTTTTGCGAAATATAACAAGAACTTAATTTC | |
| Ptet_tetR_right | TATTTTTTTTAGTTTTTCATGAACTCGAGGGG | Amplify pIMAY to insert *hsdR* upstream homology arm |
| as_secY_left | TTGACACTCTATCATTGATAGAGTATAATTAAAATAAGCTTG | |
| pIMAY_tetR_left | AGACCCACTTTCACATTTAAG | Amplify pIMAY to insert *hsdR* downstream homology arm |
| pIMAY_tetR_right | TTAAAAGCAGCATAACCTTTTTCC | |
| Ptet_minus10_CATAAT_forward | CTCTATCAATGATAGAGTGTCAATATTTTTTTTAGTTTTTC | Mutate $P_{tet}$ minus 10 to CATAAT |
| Ptet_minus10_CATAAT_reverse | CATAATTAAAATAAGCTCTCTATCATTGATAGAGTACGATC | |
| tetR_minus10_TATAAT_forward, | TAATGTCAACAAAAAGGAGGAATTAATG | Mutate *tetR* minus 10 to TATAAT |
| tetR_minus10_TATAAT_reverse | TATCATTGATAGAGTTATTTGTCAAACTAG | |
| NE1357_check_forward | TCAGACTGTGTGAGCATGGA | Confirm transposon insertion by PCR |
| NE1357_check_reverse | AGCAACATCAAGACCTCAGGA | |
| NE1420_check_forward | TGTTGGGTGATGTTATTAGTGCT | Confirm transposon insertion by PCR |
| NE1420_check_reverse | GCGCAACCATTTCAACTTTATCT | |
| 1106_us_hsdR_forward | CTCGTATTGGCTTAAATGTCAG | Confirm allelic exchange of *hsdR::dcas9-tetR* |
| hsdR_ds_1076_reverse | GTTTCCCACGCAAATAAGGA | |
| R1_no_aTc | TCCCTACACGACGCTCTTCCGATCTNNNNNNTCCCTAAAGATCTTTGACAGCTAGC | Pilot scale: generate amplicon from samples without aTc |
| R2_no_aTc | GTTCAGACGTGTGCTCTTCCGATCTNNNNNNCTTACTCAGGTTGACGCTCATCT | |
| R1_aTc | TCCCTACACGACGCTCTTCCGATCTNNNNNNGAACTAAAGATCTTTGACAGCTAGC | Pilot scale: generate amplicon from samples with aTc |
| R2_aTc | GTTCAGACGTGTGCTCTTCCGATCTNNNNNNAGGACTCAGGTTGACGCTCATCT | |
| R1v3_0 | TCCCTACACGACGCTCTTCCGATCTCAGCTAGCTCAGTCCTAGGT | Genome scale: generate amplicon from samples with or without aTc |
| R1v3_1 | TCCCTACACGACGCTCTTCCGATCTNCAGCTAGCTCAGTCCTAGGT | |
| R1v3_2 | TCCCTACACGACGCTCTTCCGATCTNNCAGCTAGCTCAGTCCTAGGT | |
| R1v3_3 | TCCCTACACGACGCTCTTCCGATCTNNNCAGCTAGCTCAGTCCTAGGT | |
| R1v3_4 | TCCCTACACGACGCTCTTCCGATCTNNNNCAGCTAGCTCAGTCCTAGGT | |
| R1v3_5 | TCCCTACACGACGCTCTTCCGATCTNNNNNCAGCTAGCTCAGTCCTAGGT | |
| R1v3_6 | TCCCTACACGACGCTCTTCCGATCTNNNNNNCAGCTAGCTCAGTCCTAGG | |

| | | |
|---|---|---|
| R2_no_aTc_0 | GTTCAGACGTGTGCTCTTCCGATCTGCTTACTCAGGTTGACGCTCATCT | Genome scale: generate amplicon from samples without aTc |
| R2_no_aTc_1 | GTTCAGACGTGTGCTCTTCCGATCTNGCTTACTCAGGTTGACGCTCATCT | |
| R2_no_aTc_2 | GTTCAGACGTGTGCTCTTCCGATCTNNGCTTACTCAGGTTGACGCTCATCT | |
| R2_no_aTc_3 | GTTCAGACGTGTGCTCTTCCGATCTNNNGCTTACTCAGGTTGACGCTCATCT | |
| R2_no_aTc_4 | GTTCAGACGTGTGCTCTTCCGATCTNNNNGCTTACTCAGGTTGACGCTCATCT | |
| R2_no_aTc_5 | GTTCAGACGTGTGCTCTTCCGATCTNNNNNGCTTACTCAGGTTGACGCTCATCT | |
| R2_no_aTc_6 | GTTCAGACGTGTGCTCTTCCGATCTNNNNNNGCTTACTCAGGTTGACGCTCATCT | |
| R2_atc_0 | GTTCAGACGTGTGCTCTTCCGATCTCAGGACTCAGGTTGACGCTCATCT | Genome scale: generate amplicon from samples with aTc |
| R2_atc_1 | GTTCAGACGTGTGCTCTTCCGATCTNCAGGACTCAGGTTGACGCTCATCT | |
| R2_atc_2 | GTTCAGACGTGTGCTCTTCCGATCTNNCAGGACTCAGGTTGACGCTCATCT | |
| R2_atc_3 | GTTCAGACGTGTGCTCTTCCGATCTNNNCAGGACTCAGGTTGACGCTCATCT | |
| R2_atc_4 | GTTCAGACGTGTGCTCTTCCGATCTNNNNCAGGACTCAGGTTGACGCTCATCT | |
| R2_atc_5 | GTTCAGACGTGTGCTCTTCCGATCTNNNNNCAGGACTCAGGTTGACGCTCATCT | |
| R2_atc_6 | GTTCAGACGTGTGCTCTTCCGATCTNNNNNNCAGGACTCAGGTTGACGCTCATCT | |
| tarO_targeting_spacer_1493_BsaI_sense | TAGTGTACCACCCATAACTGAAAT | Assemble *tarO* targeting sgRNA at the pSD14 BsaI site. |
| tarO_targeting_spacer_1493_BbsI_sense | GAAAGTACCACCCATAACTGAAAT | Assemble *tarO* targeting sgRNA at the pSD14 BbsI site. |
| tarO_targeting_spacer_1493_antisense | AAACATTTCAGTTATGGGTGGTAC | Assemble *tarO* targeting sgRNA at the pSD14 BsaI or BbsI site. |
| ltaS_targeting_spacer_1433_BsaI_sense | TAGTTGATACCACCAGATTTACCA | Assemble *ltaS* targeting sgRNA at the pSD14 BsaI site. |
| ltaS_targeting_spacer_1433_BbsI_sense | GAAATGATACCACCAGATTTACCA | Assemble *ltaS* targeting sgRNA at the pSD14 BbsI site. |
| ltaS_targeting_spacer_1433_antisense | AAACTGGTAAATCTGGTGGTATCA | Assemble *ltaS* targeting sgRNA at the pSD14 BsaI or BbsI site. |
| tarG_targeting_spacer_1277_BsaI_sense | TAGTTGATGTTAATAACGTCACCG | Assemble *tarG* targeting sgRNA at the pSD14 BsaI site. |
| tarG_targeting_spacer_1277_BbsI_sense | GAAATGATGTTAATAACGTCACCG | Assemble *ltaS* targeting sgRNA at the pSD14 BbsI site. |
| tarG_targeting_spacer_1277_antisense | AAACCGGTGACGTTATTAACATCA | Assemble *ltaS* targeting sgRNA at the pSD14 BsaI or BbsI site. |
| accC_targeting_spacer_2975_BsaI_sense | TAGTATAATCGCTTCATCTCGTGT | Assemble *accC* targeting sgRNA at the pSD14 BsaI site. |
| accC_targeting_spacer_2975_antisense | AAACACACGAGATGAAGCGATTAT | Assemble *accC* targeting sgRNA at the pSD14 BsaI or BbsI site. |
| ftsW_targeting_spacer_2050_BbsI_sense | GAAATTATGACATATGCTAATTGT | Assemble *ftsW* targeting sgRNA at the pSD14 BbsI site. |

| | | |
|---|---|---|
| ftsW_targeting_spacer_2050_antisense | AAACACAATTAGCATATGTCATAA | Assemble *fstW* targeting sgRNA at the pSD14 BsaI or BbsI site. |
| pbpA_targeting_spacer_2177_BbsI_sense | GAAAATAACTTTAGGATTTTTCTT | Assemble *pbpA* targeting sgRNA at the pSD14 BbsI site. |
| pbpA_targeting_spacer_2177_antisense | AAACAAGAAAAATCCTAAAGTTAT | Assemble *pbpA* targeting sgRNA at the pSD14 BsaI or BbsI site. |
| auxB_targeting_spacer_2031_BbsI_sense | GAAAGCAATAATAAAGAATAATAT | Assemble *auxB* targeting sgRNA at the pSD14 BbsI site. |
| auxB_targeting_spacer_2031_antisense | AAACATATTATTCTTTATTATTGC | Assemble *auxB* targeting sgRNA at the pSD14 BsaI or BbsI site. |
| gpsB_targeting_spacer_2707_BbsI_sense | GAAAGAAAAACTTTTATTGTCCTG | Assemble *gpsB* targeting sgRNA at the pSD14 BbsI site. |
| gpsB_targeting_spacer_2707_antisense | AAACCAGGACAATAAAAGTTTTTC | Assemble *gpsB* targeting sgRNA at the pSD14 BsaI or BbsI site. |
| dltX_targeting_spacer_1700_BsaI_sense | TAGTGCTTCAACATATTTATTAGG | Assemble *dltX* targeting sgRNA at the pSD14 BsaI site. |
| dltX_targeting_spacer_1700_antisense | AAACCCTAATAAATATGTTGAAGC | Assemble *dltX* targeting sgRNA at the pSD14 BsaI or BbsI site. |
| dltA_targeting_spacer_1702_BsaI_sense | TAGTGTACCGCCTGATGCTAAACA | Assemble *dltA* targeting sgRNA at the pSD14 BsaI site. |
| dltA_targeting_spacer_1702_antisense | AAACTGTTTAGCATCAGGCGGTAC | Assemble *dltA* targeting sgRNA at the pSD14 BsaI or BbsI site. |
| dltB_targeting_spacer_1705_BsaI_sense | TAGTAAAGGTTTATCGAAGTTTGG | Assemble *dltB* targeting sgRNA at the pSD14 BsaI site. |
| dltB_targeting_spacer_1705_antisense | AAACCCAAACTTCGATAAACCTTT | Assemble *dltB* targeting sgRNA at the pSD14 BsaI or BbsI site. |
| dltC_targeting_spacer_1706_BsaI_sense | TAGTCTAATAATAATCCAACTGTT | Assemble *dltC* targeting sgRNA at the pSD14 BsaI site. |
| dltC_targeting_spacer_1706_antisense | AAACAACAGTTGGATTATTATTAG | Assemble *dltC* targeting sgRNA at the pSD14 BsaI or BbsI site. |
| dltD_targeting_spacer_1708_BsaI_sense | TAGTAATCCTGTAAACCAACTAGC | Assemble *dltD* targeting sgRNA at the pSD14 BsaI site. |
| dltD_targeting_spacer_1708_antisense | AAACGCTAGTTGGTTTACAGGATT | Assemble *dltD* targeting sgRNA at the pSD14 BsaI or BbsI site. |

**Supplementary Table 6. pSD14 plasmids used for gene silencing in clonal populations.**

| Plasmid Name | BsaI spacer sequence | BbsI spacer sequence |
|---|---|---|
| pSD14 | | |
| pSD14 BsaI-*tarO* | GTACCACCCATAACTGAAAT | |
| pSD14 BbsI-*tarO* | | GTACCACCCATAACTGAAAT |
| pSD14 BsaI-*ltaS* | TGATACCACCAGATTTACCA | |
| pSD14 BbsI-*ltaS* | | TGATACCACCAGATTTACCA |
| pSD14 BsaI-*tarO* BbsI-*ltaS* | GTACCACCCATAACTGAAAT | TGATACCACCAGATTTACCA |
| pSD14 BsaI-*ltaS* BbsI-*tarO* | TGATACCACCAGATTTACCA | GTACCACCCATAACTGAAAT |
| pSD14 BsaI-*tarG* | TGATGTTAATAACGTCACCG | |
| pSD14 BbsI-*tarG* | | TGATGTTAATAACGTCACCG |
| pSD14 BsaI-*tarO* BbsI-*tarG* | GTACCACCCATAACTGAAAT | TGATGTTAATAACGTCACCG |
| pSD14 BsaI-*tarG* BbsI-*tarO* | TGATGTTAATAACGTCACCG | GTACCACCCATAACTGAAAT |
| pSD14 BsaI-*accC* | ATAATCGCTTCATCTCGTGT | |
| pSD14 BbsI-*ftsW* | | TTATGACATATGCTAATTGT |
| pSD14 BsaI-*accC* BbsI-*ftsW* | ATAATCGCTTCATCTCGTGT | TTATGACATATGCTAATTGT |
| pSD14 BbsI-*pbpA* | | ATAACTTTAGGATTTTTCTT |
| pSD14 BsaI-*accC* BbsI-*pbpA* | ATAATCGCTTCATCTCGTGT | ATAACTTTAGGATTTTTCTT |
| pSD14 BbsI-*auxB* | | GCAATAATAAAGAATAATAT |
| pSD14 BbsI-*gpsB* | | GAAAAACTTTTATTGTCCTG |
| pSD14 BsaI-*dltX* | GCTTCAACATATTTATTAGG | |
| pSD14 BsaI-*dltA* | GTACCGCCTGATGCTAAACA | |
| pSD14 BsaI-*dltB* | AAAGGTTTATCGAAGTTTGG | |
| pSD14 BsaI-*dltC* | CTAATAATAATCCAACTGTT | |
| pSD14 BsaI-*dltD* | AATCCTGTAAACCAACTAGC | |
| pSD14 BsaI-*dltX* BbsI-*auxB* | GCTTCAACATATTTATTAGG | GCAATAATAAAGAATAATAT |
| pSD14 BsaI-*dltA* BbsI-*auxB* | GTACCGCCTGATGCTAAACA | GCAATAATAAAGAATAATAT |
| pSD14 BsaI-*dltB* BbsI-*auxB* | AAAGGTTTATCGAAGTTTGG | GCAATAATAAAGAATAATAT |
| pSD14 BsaI-*dltC* BbsI-*auxB* | CTAATAATAATCCAACTGTT | GCAATAATAAAGAATAATAT |
| pSD14 BsaI-*dltD* BbsI-*auxB* | AATCCTGTAAACCAACTAGC | GCAATAATAAAGAATAATAT |
| pSD14 BsaI-*dltX* BbsI-*gpsB* | GCTTCAACATATTTATTAGG | GAAAAACTTTTATTGTCCTG |
| pSD14 BsaI-*dltA* BbsI-*gpsB* | GTACCGCCTGATGCTAAACA | GAAAAACTTTTATTGTCCTG |
| pSD14 BsaI-*dltB* BbsI-*gpsB* | AAAGGTTTATCGAAGTTTGG | GAAAAACTTTTATTGTCCTG |
| pSD14 BsaI-*dltC* BbsI-*gpsB* | CTAATAATAATCCAACTGTT | GAAAAACTTTTATTGTCCTG |
| pSD14 BsaI-*dltD* BbsI-*gpsB* | AATCCTGTAAACCAACTAGC | GAAAAACTTTTATTGTCCTG |

#### Supplementary References


1. Reed P, Sorg M, Alwardt D, Serra L, Veiga H, Schäper S, Pinho MG. 2024. A CRISPRi-based genetic resource to study essential Staphylococcus aureus genes. mBio 15:e0277323.

2. Fey PD, Endres JL, Yajjala VK, Widhelm TJ, Boissy RJ, Bose JL, Bayles KW. 2013. A genetic resource for rapid and comprehensive phenotype screening of nonessential Staphylococcus aureus genes. mBio 4:e00537-12.

3. Monk IR, Shah IM, Xu M, Tan MW, Foster TJ. 2012. Transforming the untransformable: application of direct transformation to manipulate genetically Staphylococcus aureus and Staphylococcus epidermidis. mBio 3.

4. Peters JM, Colavin A, Shi H, Czarny TL, Larson MH, Wong S, Hawkins JS, Lu CHS, Koo B-M, Marta E, Shiver AL, Whitehead EH, Weissman JS, Brown ED, Qi LS, Huang KC, Gross CA. 2016. A Comprehensive, CRISPR-based Functional Analysis of Essential Genes in Bacteria. Cell 165:1493–1506.

5. Brückner R, Wagner E, Götz F. 1993. Characterization of a sucrase gene from Staphylococcus xylosus. J Bacteriol 175:851–857.
